# Profiling Bile Acids in the Stools of Humans and Animal Models of Cystic Fibrosis

**DOI:** 10.1101/2025.05.08.651222

**Authors:** Melissa M. Carmichael, Rebecca A. Valls, Shannon Soucy, Julie Sanville, Juliette Madan, Sarvesh V. Surve, Mark S. Sundrud, George A. O’Toole

## Abstract

Cystic fibrosis (CF) is associated with dysbiosis of the gut microbiome, alterations in intestinal mucus production, aberrant bile acid (BA) metabolism, fat malabsorption, and chronic inflammation. As little is known about BAs in CF, we performed both comprehensive and targeted BA profiling in stool of children with or without CF. Our results reveal that select BA species and metabolites are significantly different between children with CF (cwCF) and healthy controls. There is also a trend towards higher primary cBA and total BA levels for cwCF. Matched bacterial metagenomic analyses showed no change in alpha-diversity between groups in our small cohort, at odds with previous studies, whereas changes in relative abundance of *Bacteroides* (lower) and *E. coli* (increased) species is consistent with prior reports. A robust trend was noted toward reduced abundance of *bsh* gene families (Wilcox test, p = 0.052), a key rate-limiting enzyme required for bacterial synthesis of secondary BAs, in cwCF. Modest changes in both BAs and microbial BA metabolism-related gene abundances may be attributable to small sample sizes, but also suggest likely combination defects in both host and microbial BA metabolic pathways in cwCF. Importantly, although fecal BA profiles from both ferret and mouse CF models showed significant differences from human BA profiles, only the ferret model reproduced significant differences between CF and nonCF animals, highlighting ferrets as a potentially more appropriate model for studying BA in stool in the context of CF. Together, these results provide new insights into CF-related BA dysmetabolism in cwCF, and highlight limitations of CF animal models for BA functional studies.

**IMPORTANCE:** Changes in the abundance and/or composition of intestinal bile acids (BAs) may contribute to dysbiosis and altered gastrointestinal physiology in CF. Here, we report shifts in select fecal BA classes and species for children with CF (cwCF). Matched metagenomic analysis suggest possible defects in both host intestinal BA absorption and gut microbial BA metabolism. Additional analyses of mouse and ferret CF stool for BA composition suggest great care must be taken when interpreting BA functional studies using these animal models. Together, this work lays technical and conceptual foundations for interrogating BA-microbe interactions in cwCF.

## INTRODUCTION

Cystic fibrosis (CF) is a hereditary disease caused by mutations in the gene encoding the cystic fibrosis transmembrane conductance regulator (CFTR) protein. Disruption of CFTR function results in alterations in the secretion of chloride and bicarbonate causing an imbalance in the hydration of mucosal surfaces, and accumulation of abnormally thick, viscous mucus in the lungs and intestinal tract (1–3). Whereas pulmonary disease dominates the adult population of persons with CF (pwCF), gastrointestinal (GI) complications are an important cause of morbidity for pwCF early in life. Infants and children with CF (cwCF) can experience an array of GI symptoms including meconium ileus, small bacterial overgrowth, and nutrient malabsorption (1, 3–6).

From an early age, cwCF exhibit a decrease in α-diversity, as evidenced by fecal 16S rRNA amplicon library and metagenomic sequencing (7–11). Literature indicates the persistence of this dysbiosis into adulthood among pwCF (3, 5, 12, 13). Notable taxonomic differences observed in cwCF include a reduction in specific immune-modulating bacterial genera in the phyla Bacteroidetes (i.e., *Bacteroides* spp.), which is positively associated with gut health, and an increase in pathogenic Proteobacteria (i.e., *Escherichia coli*) (7–11). These shifts in colonic microbial communities are likely driven by changes in the physiological features of the intestinal environment (2, 14). The physiological features of the CF colon can be characterized by excess mucus and fat content, acidic pH, inflammation, antibiotic perturbations as well as altered bile acids (BA) (15–18).

BAs play key roles in shaping gut microbial colonization and function, acting as detergents and supporting the absorption of fats by intestinal enterocytes (19–21). While the liver is responsible for the production of primary BAs (pBAs), those that escape intestinal absorption in the distal small intestine (i.e., ileum) become increasingly subject to modification by microbiota in the large intestine. The products of bacterial BA metabolism (i.e., unconjugated BA and secondary BA species and metabolites) dramatically increases the structural and functional diversity of BAs in enterohepatic circulation (19, 21). BAs have direct anti-microbial functions, impacting susceptible bacteria in both a bacteriostatic and bactericidal fashion, via disruption of bacterial membranes (20, 21). Consequently, the modification of BAs is an essential microbial defense mechanism (20, 21). For example, when tested against *Lactobacillus* and *Bifidobacterium*, pBAs were shown to disrupt membranes in a dose-dependent manner, while unconjugated BAs exert a greater reduction in viability than their conjugated counterparts (19, 22).

Among the hallmarks of GI complications in pwCF, as well as in murine models, is a reported 3-fold increase in fecal BA excretion (20, 21, 23–25). Enterohepatic circulation of BAs is a tightly regulated system in which ∼95% of total BAs are reabsorbed in the ileum and the remaining ∼5% pass through the colon and are excreted via the feces (21). In a healthy individual, the secondary BAs lithocholic acid (LCA) and deoxycholic acid (DCA) dominate fecal profiles (26). Previous studies suggest that impaired BA homeostasis manifest as BA malabsorption and an increased cholic acid (CA) to chenodeoxycholic acid (CDCA) ratio among duodenal BAs in adult pwCF (20, 23, 24). Two studies have been performed using samples from children ages ranging from 2 years to 20 years (25, 27). However, no studies have focused on BA homeostasis in cwCF from birth to 36 months of age, the key window for immune programming (28, 29), and on the relationship between BAs and gut microbial communities in the CF gut. Previous studies also profiled only a fraction – typically a small handful – of the estimated hundreds of BA species now recognized in the human intestinal tract. We address these knowledge gaps here.

Improved animal models have emerged as key tools for deciphering mechanistic links between gut microbiota and disease in CF (2, 30). Thus, we also sought to examine the translational relevance of BA profiles within two common and notable CF animal models – ferret and mouse. We show that stool BA profiles in ferrets and mice with CF do not parallel those of cwCF, although there is a difference in BA profiles in ferrets with and without CF. Hence, caution must be taken when selecting animal models for the study of BA functions in the context of CF. Altogether, these data provide key new insights for understanding relationships between BAs and microbial communities in the gut of cwCF, as well as for developing translationally relevant in vitro and in vivo model systems to disentangle complex relationships between intestinal BAs and microbiota. These investigations also lay the groundwork for clairfying interventions leveraging the complex micro- and macro-environment of the intestine in cwCF with the goal of optimizing systemic health.

## RESULTS

### CF-related alterations in the fecal bile acid metabolome amongst children

Using cutting edge metabolomics, we surveyed 89 BA species and related metabolites in feces (stool) from individual children with cystic fibrosis (cwCF) and healthy controls (**Figure S1A**). Fecal samples capture the fraction of BAs that escape ileal absorption and are metabolized in the large intestine. Liquid chromatography with tandem mass spectrometry (LC-MS/MS) quantified 89 discrete BA species and related metabolites. 84 of 89 species examined displayed values >0 in at least one of the samples (CF or nonCF) and were thus considered detectable. These BA species and metabolites clustered into 7 general classes based on biosynthetic origins (**Figure S1B**): (i) primary conjugated BAs (primary cBA), which are BAs synthesized and secreted by the liver at homeostasis; (ii) primary unconjugated BA (primary uBA), which are products of bacterial primary cBA deconjugation; (iii) secondary conjugated BAs (secondary cBA), which are secondary uBAs re-conjugated in hepatocytes; (iv) secondary unconjugated BA (secondary uBA), which are products generated from primary uBAs via bacterial metabolism; (v) secondary metabolites, which are more extensively modified secondary uBAs produced by bacterial metabolism; (vi) hepatic detoxification products, including sulfate and glucuronide conjugates produced to limit BA toxicity and promote clearance; and (vii) synthetic, intermediates, and atypical BAs, which are normally secreted in trace amounts by healthy livers, but can increase during active liver disease (31).

Comparison of log transformed total concentration for each of the seven BA classes between CF and nonCF showed BAs derived from hepatocytes – primary cBAs, primary uBAs, hepatic detoxification products and synthetic, intermediates, and atypical BAs – were elevated in stool of cwCF vs. controls, whereas those produced directly via microbial metabolism (<u>e.g</u>., secondary uBAs and metabolites) were reduced (**Figure 1A**). Total BA levels between cwCF versus controls showed no statistically significant difference (**Figure 1B**) – contrary to previous findings in adults with CF (20, 21, 23–25) – although there was a trend towards higher total BA level in cwCF, which does parallel previous studies (**Figure 1B**). CF-associated changes in fecal BA pool composition, more so than in overall levels (**Figure 1C, Table S1, Figure S2A-B**), suggest that aberrant BA metabolism begins manifesting early in cwCF. Further, we used a linear model for each of the individual BAs to determine significant differences in log2 transformed concentrations (**Figure S3-8, Table S2**), accounting for genotype (CF, nonCF) and batch of samples analyzed (batch 1 or batch 2). Of the 84 BAs detected, only alloisolithocholic acid, taurolithocholic acid 3 S, deoxycholic acid 3 G and lithocholic acid 3 G have an adjusted p-value <0.05 (**Table S2**), and thus a statistically significantly difference in the CF versus nonCF cohort.

**Figure 1.**
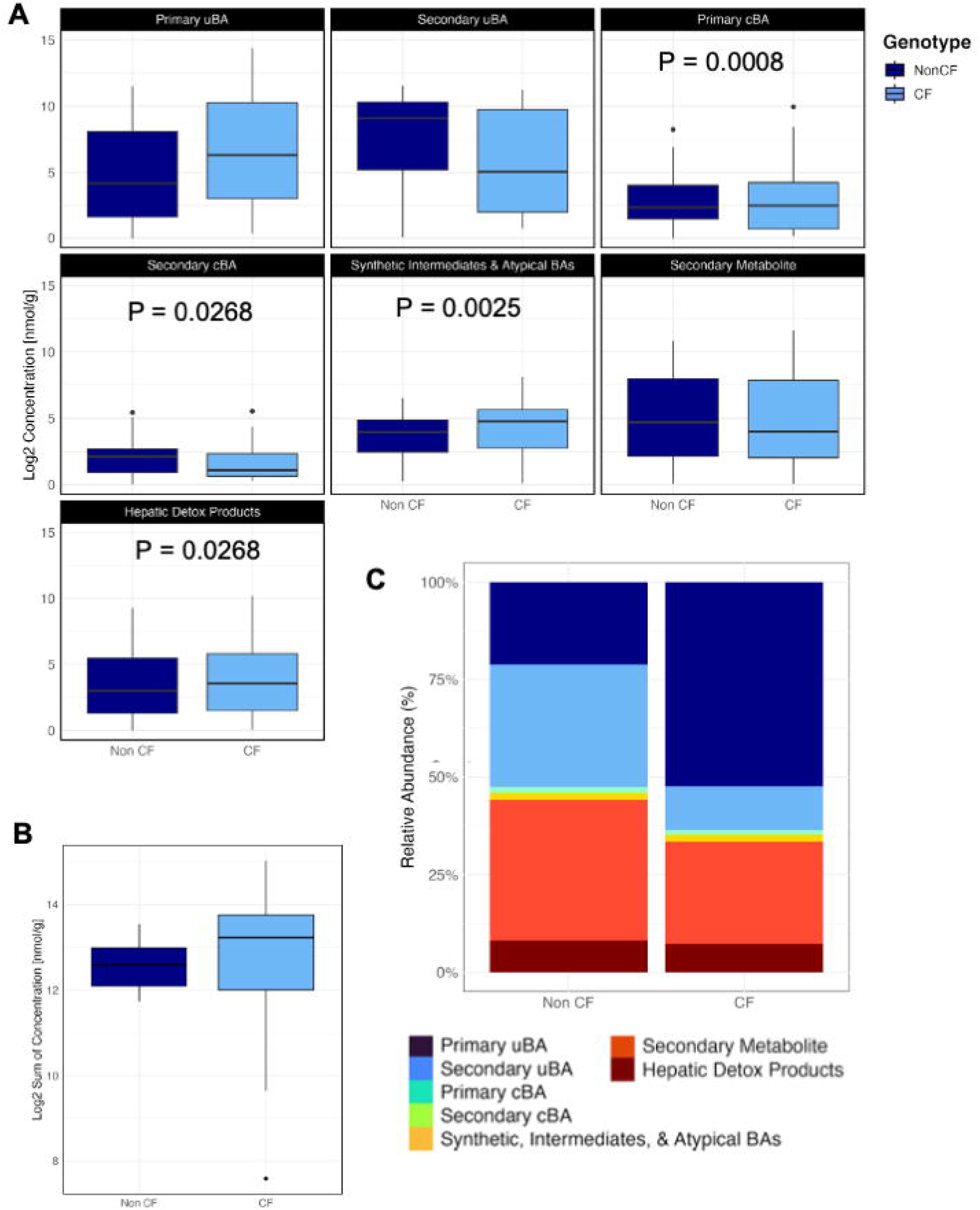
Summary of untargeted BA profiles from human stool samples. **A.** Box plots comparing log2 transformed concentration for each of the seven classes from human fecal samples (n =15) of each genotype (CF and nonCF). Statistical significance was determined by linear model, which is summarized in **Table S1-2**. **B**. Total bile acid concentration (log2 transformed) for each sample from the same human fecal samples in Panel A (n =15) of each genotype (CF and nonCF). Linear model does not show significance (P-value = 0.929). **C.** Relative abundance of functional groups between genotypes (CF and nonCF) in human samples.

We identified the BA species most differentially abundant between cwCF and non-CF controls by calculating the difference in mean log2 concentrations for each bile acid between genotypes. To visualize these shifts, we plotted the log2-transformed concentrations of each bile acid species from cwCF and non-CF stool samples against their respective genotype averages, highlighting the species with the greatest differences (**Figure S2A**). Using this analysis, the top differentially abundant BA species for both genotypes were selected to develop a targeted panel for subsequent studies (**Table S3**).

To explore BA species further, we quantified a subset of the 21 most differentially abundant BA species, including known bacterial metabolites (**Table S3**), in eighteen CF and eighteen nonCF, which included the 30 fecal samples analyzed in **Figure 1** plus 3 additional samples for each genotype. Comparison of log2 transformed concentration of BA classes showed a significant increase for two of the classes in cwCF: primary uBA as well as synthetic, intermediates, and atypical BAs (**Figure 2A, Table S4**). Total BA excretion of the targeted BA panel shows a significant increase in BA levels in CF samples compared to healthy controls (**Figure 2B**), as has been seen in adult populations (20). We noted marked differences in BA between CF and non-CF samples (**Figure 2C, Figure S2C**). Amongst individual BA species, linear modeling with corrections for multiple comparisons indicates that only beta muricholic acid and allocholic acid are different between groups after adjusting for multiple comparisons (**Figure S9-10, Table S5**). Enrichment of unconjugated primary BAs, together with the overall increase in fecal BA excretion is consistent with both BA and fat malabsorption noted in CF.

**Figure 2.**
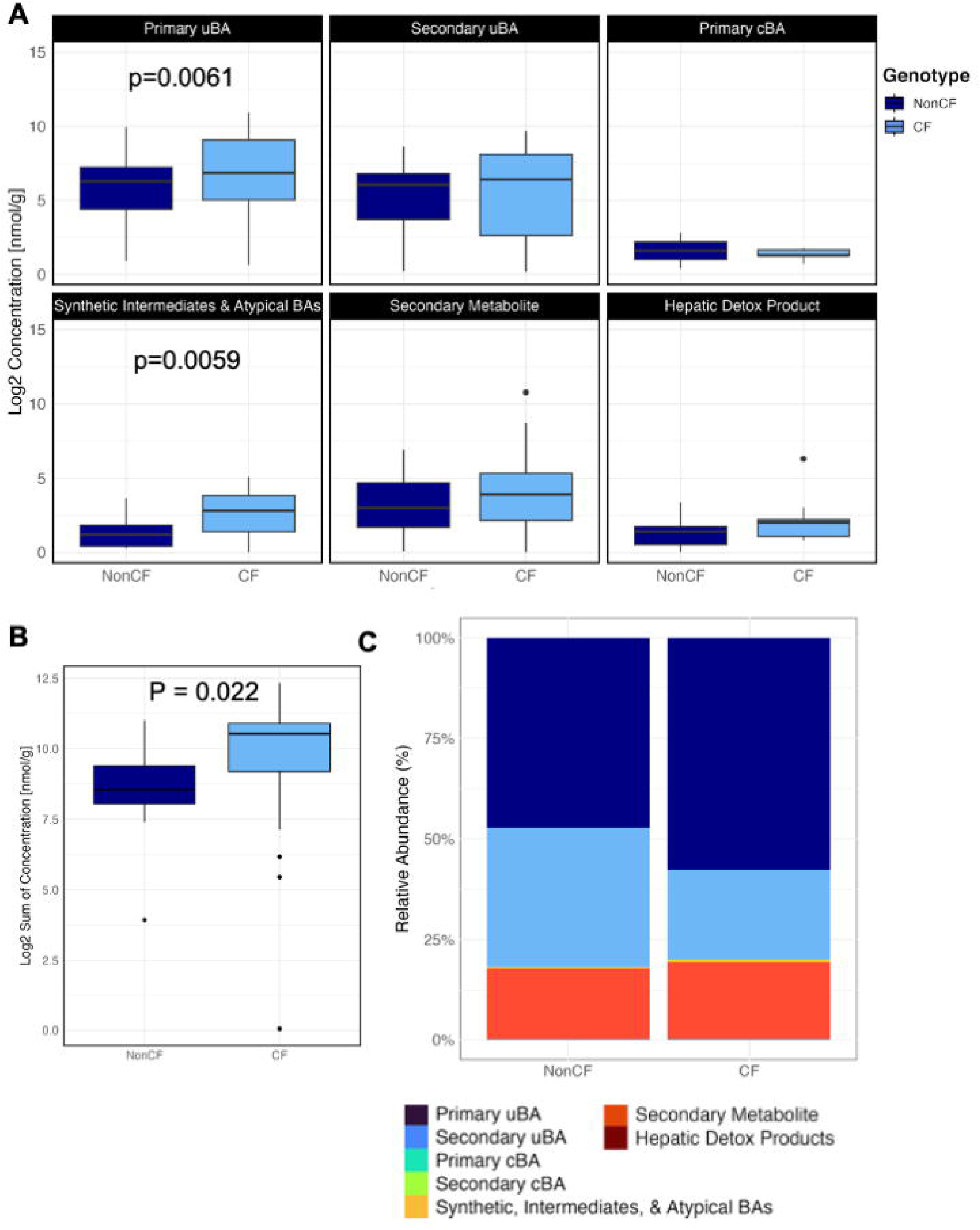
Summary of targeted (most abundant and differentially abundant) BA profiles from human stool samples. **A**. Box plots comparing log2 transformed concentration for each of the seven functional groups from human fecal samples (n = 18) for each genotype (CF and nonCF). Data analysis performed by linear model, which is summarized in **Table S4-5**. **B**. Total bile acid concentration (log2 transformed) for each sample from the same human fecal samples in Panel A (n =18) of each genotype (CF and nonCF). A Wilcoxon rank-sum test, consistent with methodologies previously applied in adult populations (20), revealed a significant difference between the sums of BA excretion between genotypes (p = 0.022). **C.** Relative abundance of functional groups between genotypes (CF and nonCF) in human samples.

We also plotted the primary uBA versus the primary conjugated BA for each CF and nonCF subject (**Figure S2D**). This analysis revealed that increases in fecal primary uBAs among cwCF correlated with commensurate elevations in primary cBA precursors, suggesting that CF-related increases in fecal BA levels generally, and fecal primary uBAs specifically, at least partly involves a combination of elevated hepatic BA synthesis and impaired ileal BA absorption.

### Metagenomic analysis of human fecal samples

To explore the microbial composition associated with the human samples, we performed shotgun metagenomic sequencing in 15 CF and 15 nonCF human fecal samples, for which we had parallel comprehensive bile acid profiles on (see **Figure 1**). α-diversity was calculated using the Shannon Diversity Index, accounting for both richness (number of species) and evenness (distribution of species abundances). Overall, no significant differences were found between nonCF and CF samples (**Figure S11**), although there was a trend towards lower α-diversity for CF samples, which is expected based on previous findings and perhaps consistent with a reduced effect sizes in young children.

Taxonomic abundances indicate sample CF4 is a clear outlier, showing that 80% of total abundance is dominated by only 2 phyla (**Figure S12**), while CF4, CF8 and CF14 show particularly high abundances of Proteobacteria (**Figure S12**). A trend in the relative abundance of *Bacteroides* (significantly reduced) was noted (**Figure S13**).

We next mined our metagenomes for abundance of BA metabolism-related genes (**Figure 3A**). We observed a strong trend towards reduced levels of *bsh* gene families within stool microbial communities of cwCF (**Figure 3B-C**), whereas a key member of the *bai* operon, *baiE,* which encodes 7-α−dehydroxylase and acts downstream of *bsh*, shows a modest and non-significant increase in the CF samples (**Figure 3D-E**). Together, these data suggest that depletion of bacterially-derived secondary BA species and metabolites in cwCF (**Figure 2**) may be attributed to lower rates of the first step in bacterial BA biotransformation, deconjugation via *bsh*. At the same time, CF-related increases in primary uBA species – that are direct products of *bsh*-mediated deconjugation and that parallel increases in primary cBA precursors (**Figure S2D**) – could be even higher in cwCF if not for dysbiosis limiting community-wide BA deconjugation machinery.

**Figure 3.**
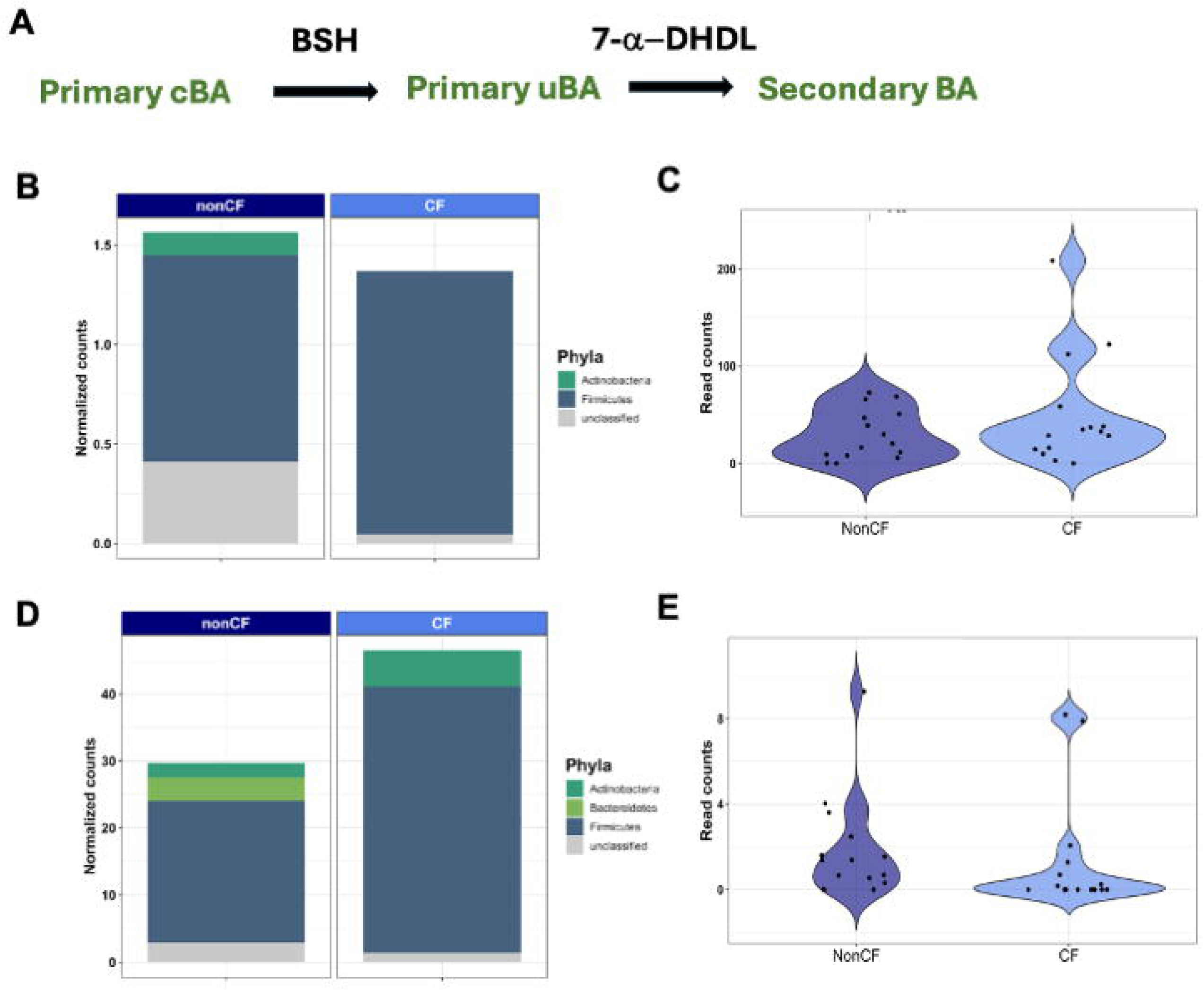
Analysis of microbial bile metabolism genes in human stool samples. **A.** Simplified bacterial BA catabolism pathway showing the role of BSH in the deconjugation of primary BA, and 7-α-dehydroxylase participating in the conversion of these molecules to secondary BA. **B.** Normalized counts of the *bsh* gene in CF and nonCF controls. Reads were mapped to the UniRef_90 database, which represents gene families clustered at 90% identity. Clustering at 90% identity enables assignment of taxonomic groups for most gene families. Counts for each gene family are first normalized for gene length with reads per kilobase and then normalized for library depth with total sum scaling to enable comparison between samples. Read counts for each gene families were averaged across samples from the same genotype (nonCF and CF) and the sum of the average read count for gene families representing bile salt hydrolase is represented for each genotype. **C.** The read counts for all *bsh* gene familes were summed for nonCF and CF samples and the distributions tested for significant different between groups using the Wilcox test (p = 0.052). **D.** Normalized counts of the *baiE* gene, which codes for the 7-a-dehydroxylase. These data were generated as described in panel B. **E**. The read counts for all gene familes encoding 7-α-dehydroxylases were summed for nonCF and CF samples and the distributions tested for significant different between groups using the Wilcox test (p = 0.081).

We next asked if CF-related shifts in BA composition is due to dysbiosis of specific microbes. Consistent with this hypothesis, circos plots indicating correlations between select BA species and phyla show clear differences in the association between phyla and BA for nonCF versus CF samples (**Figure S14A-B**). For example, for CF samples there is a negative correlation between Bacteroidetes and the primary cBA, cholic acid, as would be expected given that this group of organisms can carry the *bsh* genes. Interestingly, for CF samples, many of the correlations between BA are with low abundance phyla that do not change significantly between CF and nonCF samples (**Figure S14C**), indicating perhaps a complex relationship between microbial community composition and BA abundance that may be additionally influenced by changes in host intestinal BA absorption.

### Characterizing mouse and ferret BA profiles to assess their utility for studying BA in CF

To determine whether or how CF-related changes in BA metabolism are reflected in common animal models, we used our same targeted BA panel (**Table S3**) to analyze fecal BAs in mouse and ferret models of CF, compared with wild type controls. Data from WT C57BL/6 mice (nonCF) and C57BL/6 *cftr*F508del (CF) showed no significant differences in either total BA levels or levels of select BA classes between CF and nonCF animals, although CF mice again showed modest trends towards increased fecal levels of both primary uBAs and synthetic intermediates, atypical BAs (**Figure 4, Figures S15-S17, Table S6-7**). Linear modeling with corrections for multiple comparisons indicates that dehydrolithocholic acid is the only individual BA species from the mouse BA pool that is significantly different in CF versus control mice (**Table S7**).

**Figure 4.**
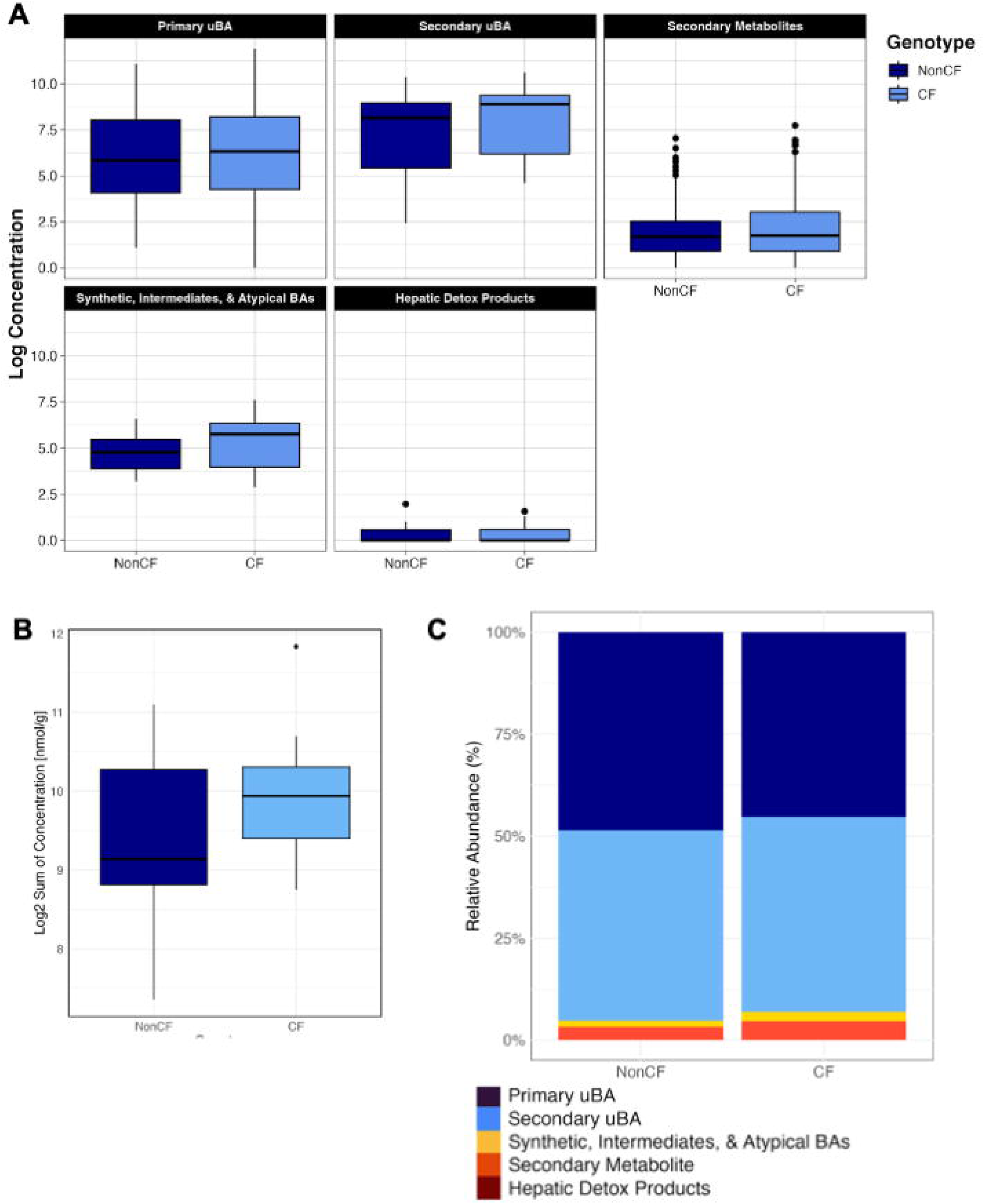
Summary of targeted BA profiles from mouse stool samples. **A**. Box plots comparing log2 transformed concentration for each of the seven functional groups from mouse fecal samples (n = 15) of each genotype (CF and nonCF). Data analysis performed by linear model, which is summarized in **Table S6-7**. **B.** Total bile acid concentration (log transformed) for each genotype from the same mouse fecal samples in Panel A (n =15) of each genotype (CF and nonCF). Linear model dose not show significance (P-value = 0.15). **C.** Relative abundance of functional groups between genotypes (CF & nonCF) in mouse samples.

Ferret samples show more statistically significant differences in CF-related fecal BA profiles, although several of these changes are paradoxical relative to our human findings. CF ferrets do show a modest but significant increase in fecal levels of primary cBA, akin to cwCF, while all other BA classes – and total fecal BA levels – are reduced compared with wild type controls. (**Figure 5A-C, Figure S18-20, Table S8**). Furthermore, measuring total BA levels shows that nonCF ferrets have significantly more BAs in their stool than ferrets carrying the mutation causing CF (**Figures 5C**), in contrast to the literature (20, 25, 27) and our human data presented here showing more BA in CF (**Figure 2B**). When investigating individual BA species, linear modeling with multiple corrections indicates 9/23 BA are significantly different in CF (**Figures S18-20, Table S9**). These data show clear differences between human data and data collected from the ferret model.

**Figure 5.**
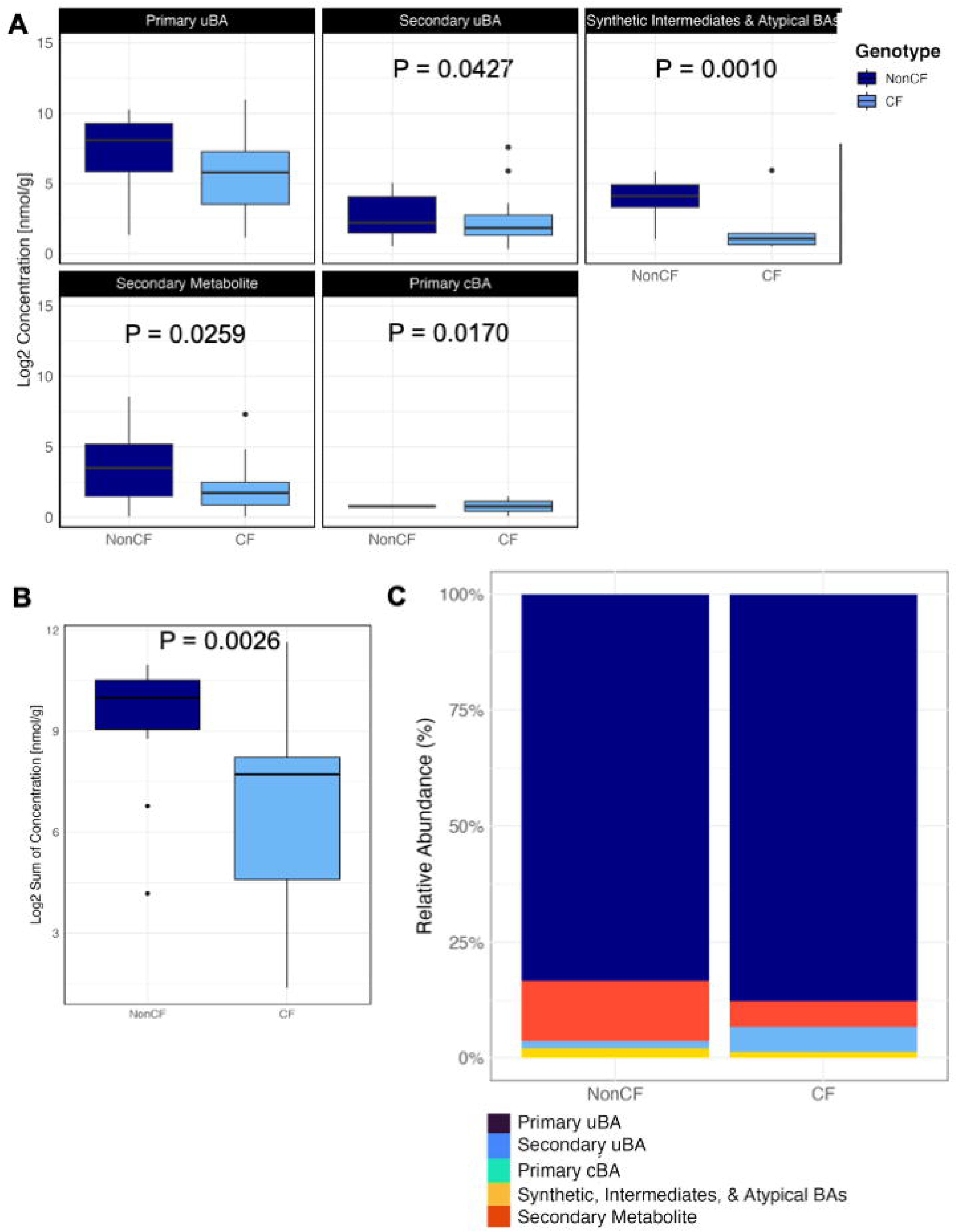
Summary of targeted BA profiles from ferret stool samples. **A**. Box plots comparing log2 transformed concentration for each of the seven functional groups from ferret fecal samples (n = 15) of each genotype (CF and nonCF). Data analysis performed by linear model, which is summarized in **Table S8 – 9**. **B.** Total bile acid concentration (log2 transformed) for each genotype from the same ferret fecal samples in Panel A (n =15) of each genotype (CF and nonCF). Linear model shows significance (P-value = 0.0026). **C.** Relative abundance of functional groups between genotypes (CF and nonCF) in ferret samples.

The observation that we saw more differences in the BA between CF and nonCF ferrets prompted us to examine the BA profiles in more detail. Analysis of the ferret BA pools showed a significant difference in the β-diversity of BA for the CF versus nonCF populations, which was also observed for the human cohort (**Figure 6A,B,D**). There is no significant difference in the β-diversity of BA for the mouse CF and nonCF samples (**Figure 6C**). **Figure 6E** shows an overlap in BA profiles for nonCF samples from humans, mice and ferrets, but there is little overlap specifically between mice and ferrets. An overlap in BA profiles for CF samples is only observed for persons and ferrets with CF. In all cases, the BA profiles are significantly different between humans and animal models.

**Figure 6:**
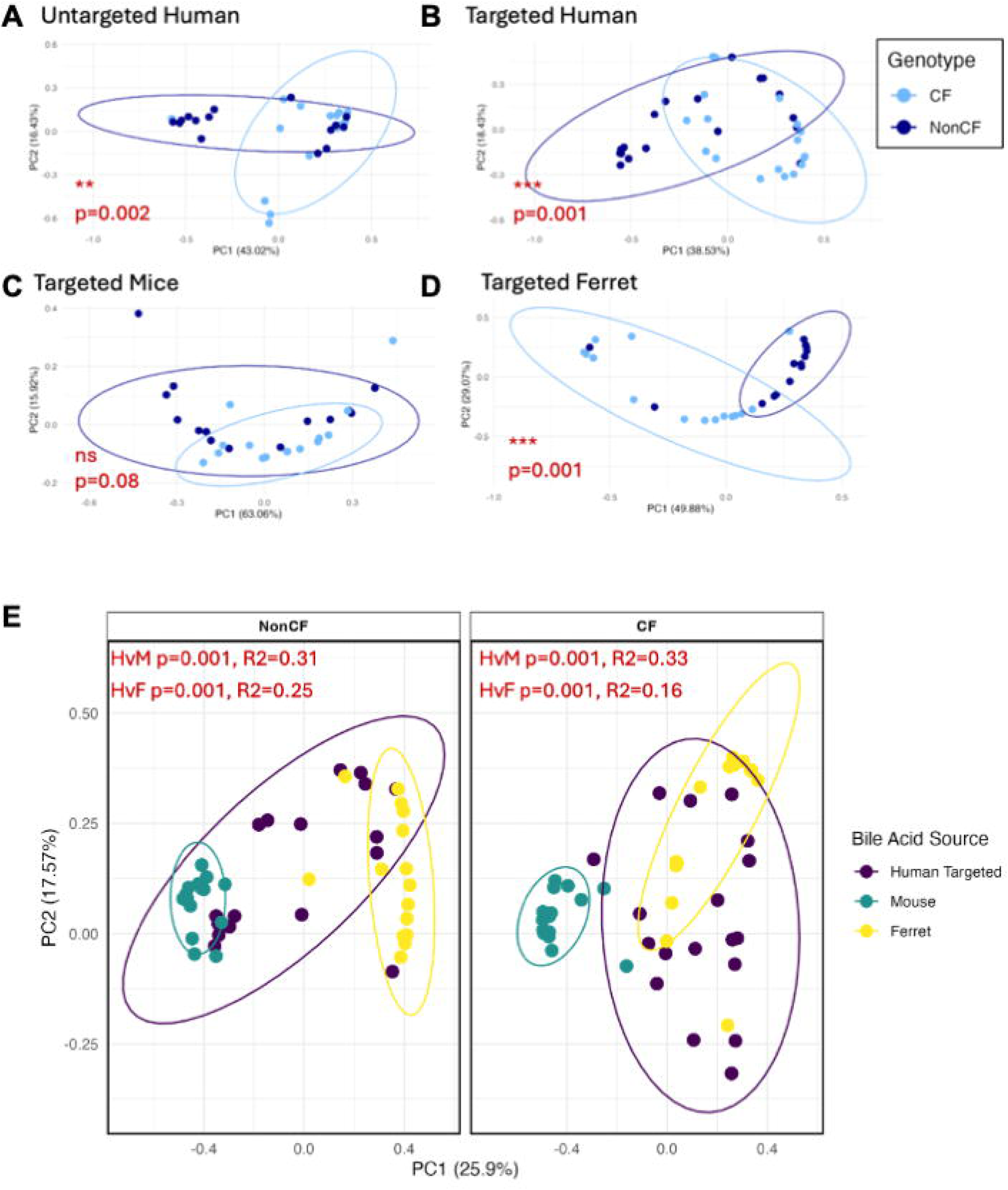
Bray Curtis beta analysis between nonCF and CF for. **A)** untargeted human bile acids, **B)** targeted panel human bile acids, **C)** targeted mouse bile acids, and **D)** targeted ferret bile acids. **E)** Bray Curtis beta analysis between bile acids sample sources for NonCF (left) and CF (right) samples. Statisical analysis was performed by PERMANOVA, where *p* and R-squared values are indicated in red. Pairwise-PERMANOVA was performed for data in panel **E**, where *p*-values indicating significance between human and mouse are indicated by HvM, and human vs ferret indicated by HvsF.

## DISCUSSION

Our findings provide a comprehensive analysis of BA profiles in fecal samples from cwCF compared to healthy controls, highlighting unique aspects of BA levels in CF. Total BA levels were significantly higher in the CF samples and distinct differences in BA composition were observed. The increase in primary uBAs underscores the potential effects of microbial dysbiosis impairing BA metabolism in pwCF, either alone or in combination with changes in host intestinal BA absorption, while differences in biosynthetic intermediates and atypical BAs for CF samples likely reflect differences in hepatic metabolism. We note that we do not observe significant differences in total fecal BA levels when analyzing the large/comprehensive panel of 89 BA, whereas we do see such differences for the targeted BA analysis. The latter observation has been what was reported previously, as described above for both CF and IBD.

Our metagenomic analysis provides data that can be mined to assess questions related to microbial BA metabolism because we can link microbial communities and genes directly to the level of BA measured given these data were acquired from the same samples. In this small cohort we did not observe statistical significance in BA-metabolic genes but trends toward a robust decrease in *bsh* abundance and a modest increase in the gene coding for 7-a-dehydroxylase (*baiE*) in CF were observed. Overall, these data also provide a resource for investigators to start a more detailed analysis of the interactions between BA, the microbiota and the host. It will be important for future studies to also incorporate meta-transcriptomics analyses, to assess relationships between BAs and not only bacterial BA metabolism-related gene abundance, but also their potential expression.

We also investigated the potential use of animal models to provide critical insights into BA metabolism in CF. Mouse models exhibit minimal differences between CF and nonCF genotypes, with only a single BA showing significantly different levels in CF versus nonCF samples. We also noted no changes in bulk levels of BA, which is at odds with the data from cwCF. It is well established that mouse livers, unlike those in humans, utilize a cytochrome P450 enzyme, Cyp2c70, to metabolize the major human pBA, CDCA, into rodent-biased muricholates (*e.g.,* αMCA, βMCA) (32, 33). MCA species are vastly more hydrophilic, and thus less potentially toxic, compared with CDCA. Hence, the utility of *cftrF508del* mice in the study of CF-related BA functions using stool sampling is unclear and should be approached with caution.

Our data show that the ferret models also display differences from humans. Ferrets show a trend towards reduction in unconjugated primary BA, while humans show a significant increase. Ferrets also show a significant reduction in secondary unconjugated BA, while humans show an increasing trend. Importantly, while pwCF show a significant increase in total BA, CF ferrets show a significant decrease. Interestingly, unlike mice, ferrets do a show a shift in BA profiles in CF versus nonCF cohorts. However, given the difference with adults and cwCF, again care must be taken when using stool samples with this model.

Beyond this work, specific implications for the distinctive bile acid pools in humans, mice and ferrets – as they relate to CF-associated BA functional studies – include the dearth of key CDCA- and microbe-derived immunoregulatory BAs, namely LCA and its metabolites, in mice (34–36). There is also the near absence of all secondary BAs (e.g., DCA and LCA) in ferrets that at least partly relate to the evolution of host BA metabolism in mammals (37). Thus, care must be taken when extrapolating findings from animal models to humans regarding BA metabolism.

We note several important limitations of our study. First, the cohort size is relatively small, constrained by the cost of the BA analysis. Second, we do not control for the diet of the human subjects, thus this limitation could cause increased variance in our data. Third, while we examined the BA profiles of cwCF here – an age range not yet assessed in the literature for both BA levels and metagenomic profiling, we profiled BA in adult animals. Notably, samples from cwCF and animal models were collected in a non-fasting state, and diet and timing of food intake could not be controlled. Given the overall similarities of BA profiles in children, teens and adults in human cohorts, our data suggest that the use of adult animals in future studies could be relevant. However, as noted above, the utility of these mouse and ferret models using stool samples for the study of BA in CF is unclear. Given the mild nature of the CF allele in our mouse model used here (38, 39), it may be necessary to explore BA functions in mouse settings that express more severe loss-of-function *Cftr* alleles.

This study provides a foundation for further exploration of the role of BA metabolism and microbial dysbiosis in CF, and provides baseline data for establishing in vitro models. Future research should focus on understanding the mechanisms underlying these changes, their impact on disease progression, and the potential for targeted therapeutic interventions to restore BA homeostasis in CF.

## MATERIALS & METHODS

### CF human cohort sample collection

Infant fecal samples were collected longitudinally from 13 days up to 7.5 yo. A total of 15 CF samples were analyzed for the large, untargeted BA panel and metagenomics. 18 CF samples were analyzed for the targeted BA panel – these samples included the same 15 samples used for the large BA panel and metagenomic study plus an additional 3 CF samples that were needed based on our power analysis (not shown). Each subject provided one sample. CF fecal samples were collected by parents and initially stored in a home freezer prior to the transfer to clinics throughout New England during routine visits with clinicians. At the clinic, the samples were stored at -80C until they were transported to the lab to be aliquoted, stored in -80C and later processed for metabolomics and sequencing. This study was approved by the Dartmouth College Committee for the Protection of Human Subjects (CPHS Study # 00021761).

### NonCF human control cohort collection

Fecal samples were collected at a local daycare center and stored briefly on ice prior to the transfer to the lab at Dartmouth. A total of 15 nonCF samples were analyzed for the large BA panel and metagenomics. 18 nonCF samples were analyzed for the targeted BA panel – these samples included the same 15 samples used for the large BA panel and metagenomic study plus an additional 3 nonCF samples that were needed based on our power analysis (not shown). At the lab, the samples were aliquoted and stored in -80C and later processed for metabolomics and sequencing. These are de-identified samples from children <3 years of age. No other additional metadata were collected. This study was approved by the Dartmouth College Committee for the Protection of Human Subjects (CPHS Study #02001623).

### CF and NonCF mouse stool collection

CF mouse fecal samples were collected from 7 female and 8 male Cftr^em1Cwr^ (an F508del with a milder CF phenotype) mice at 9 weeks; we have used these animals in our previous studies (30). NonCF mouse fecal samples were collected from 7 male and 8 female WT C57BL/6 mice at 7 weeks. Studies requiring mice for collection of stool samples were performed in accordance with a protocol that adhered to the *Guide for the Care and Use of Laboratory Animals* of the National Institutes of Health (NIH) and under the supervision of the Institutional Animal Care and Use Committee at Dartmouth College (approval #00002184). The Dartmouth College animal program is registered with the U.S. Department of Agriculture through certificate number 12-R-0001, operates in accordance with Animal Welfare Assurance (NIH/PHS) number D16-00166 (A3259-01). The program is accredited with the Association for Assessment and Accreditation of Laboratory Animal Care International (accreditation #398). Age-matched and sex-matched animals were used. Note that these was no fasting before collection of samples.

### CF and NonCF ferret stool collection

Ferret fecal samples were provided by J. Engelhardt at the University of Iowa. Ferrets were raised as described previously (40). Samples were collected, aliquoted, and stored in -80°C prior to shipping to Dartmouth. CF ferret samples included 8 females and 7 males, ages ranging from 6 to 30 months. CF ferrets were off VX-770 for an average of 133 days. NonCF ferret samples included 6 females and 9 males, ages ranging from 6 to 34 months. These studies were approved by the University of Iowa Institutional Animal Care and Use Committee. The list of animals used in this study and associated metadata is shown in **Table S9**. Note that these was no fasting before collection of samples.

### Metabolite quantification and analysis (Creative Proteomics)

Stool samples were stored frozen at -80 degrees prior to metabolite quantification. Thirty human fecal samples (100mg/sample; 15 CF and 15 nonCF) were shipped frozen to Creative Proteomics for bile acid quantification by Ultra Performance Liquid Chromatography Mass Spectrometry (UPLC-MS/MS). Samples were lyophilized to dryness, weighed and added to a 2mL homogenizing tube. The samples were homogenized in 20 uL of a 75% acetonitrile solution per mg of biomass on a MM 400 mill mixer, at 30 Hz for 3 min, followed by sonication in an ice-water bath from 5 min. Samples were centrifuged to remove debris. Clear supernatant of each sample was diluted 10-fold with internal standard (IS) solution. 5 uL aliquots of the resultant sample solutions of each of the calibrated solutions were resolved via UPLC-MS/MS.

An Agilent 1290 UHPLC system coupled to an Agilent 6495 QQQ mass spectrometer was used to analyze the BA profiles. The MS instrument was operated in the multiple reaction monitoring (MRM) mode with negative ion detection. A Waters C18 column (2.1*150 mm, 1.7 uM) was used for LC separation and the mobile phase was 0.01% formic acid in water and in acetonitrile for binary—solvent gradient elution. A mixture of standard substances of all targeted bile acids at 10uM for each compound was prepared in an IS solution of 14 deuterium-labeled bile acids. This solution was further diluted step by step to have 9-point calibration solutions.

Concentrations of detected bile acids were calculated by interpolating the internal standard calibrated linear-regression curves of individual bile acids, with the analyte-to-IS peak area ratios measured from injections of the sample solutions. The full data set is shown in **Table S10**.

### Targeted metabolite quantification and analysis

A targeted subset of BA (**Table S3**) was performed using LC-MS. Stool samples were stored frozen at -80C prior to metabolite quantification. Thirty-six human fecal samples (18/ per genotype) were thawed on ice for 30 min. Stool was weighed (<10mg) in an Eppendorf tube, 1mL of extraction buffer (0.2g butylated hydroxytoluene in 2mL milliQ-water) was added to the stool sample. Samples were vortexed for 1 minute, placed on ice, centrifuged at 10,000g at 4C for 10 minutes. Following centrifugation 750uL of the supernatant was transferred to a 2mL glass vial to be shipped to Michigan State University Mass Spectrometry and Metabolomics Core for analysis. The full data set is shown in **Table S11**.

### Description of statistical analysis for BA

Concentrations of BA are normalized by stool weight then log_2_ transformed. Any BA that was below the limit of detection in all samples (both CF and nonCF) were removed from further analysis. When samples had a concentration of 0 in a specific bile acid, a value of 1 was added to the last decimal place so that normalization (log2 transformation) across all samples could be conducted. In the untargeted BA profiling, we could detect 84 out of 89 BAs in at least one sample. Those left out were (Chenodeoxycholic acid 3G, Lithocholic acid 24 G, Hyodeoxycholic acid 3 glucuronide, Hyodeoxycholic acid 24 glucuronide, and Ursodeoxycholic axid 3 glucuronide. In order to determine statistical differences by genotype (nonCF or CF) on bile acid concentrations, we ran simple and mixed effects linear models on each individual bile acid and each bile acid functional group, respectively. We set nonCF as the reference, and set log2 transformed concentrations of BAs as the independent variable, accounting for genotype and batch (1 or 2) as the dependent variables. For bile acid functional group, we also take into account repeated measures from samples by setting sample (or human participant, mouse, or ferret) as a random effect. P-values were then adjusted for multiple comparisons (84) by False Discovery Rate (FDR) at a threshold of 0.05. To assess differences in bile acid composition between sample sources, we performed Principal Coordinates Analysis (PCoA). Briefly, we calculated a Bray-Curtis β-diversity dissimilarity matrix and visualized it using PCoA. To assess statistical significance, we ran pairwise PERMANOVA tests within each genotype comparing human versus mouse samples, and human versus ferret samples. To evaluate the relationship between primary conjugated bile acids (cBA) and primary unconjugated bile acids (uBA), bile acid concentrations were summed per sample by functional group for each metabolomics method (untargeted and targeted) in human samples. Statistical analyses, including Spearman correlation tests, were conducted to assess associations between primary cBA and primary uBA concentrations by genotype and metabolomics method. The coefficient of determination (R²) and p-values were reported to evaluate the strength and significance of these relationships. All statistical analyses were performed in R (version 4.4.2). Data processing, visualization, and statistical analysis were performed in R (v4.4.2) using the packages readr for data import, dplyr and tidyr for data manipulation, ggplot2 for plotting, lme4 and lmerTest for linear mixed-effects modeling, and vegan for diversity analyses and PERMANOVA testing.

### Shotgun metagenomic sequencing

DNA for metagenomics analysis was quantified by Qubit (Thermo Fisher). 150ng DNA was used as input into the Illumina DNA Prep kit (Illumina) for library preparation following manufacturer’s instructions. Libraries were pooled for sequencing on a NextSeq2000 instrument targeting 30M, paired-end 150bp reads per sample. The data are available under BioProject PRJNA1244851.

### Metagenomic analysis

To assess microbial diversity metagenomic data was analyzed using the standard metagenomics workflow used by the Dartmouth Genomic Data Science Core. Briefly, this workflow involves filtering host reads using KneadData v. 0.12.0, taxonomic assignment using metaphlan v. 4.0.6 with the mpa_vOct22_CHOCOPhlAnSGB_202212 database, gene family and pathway assignment with humann v. 3.7 with the full_chocophlan.v201901_v31 and uniref90_annotated_v201901b_full databases, and lastly differential abundance testing across conditions of interest with MaAslin v. 1.14.1 (41–44). All code is available upon request.

## ACKNOWLEDGEMENTS

We thank the University of Iowa Ferret National Resource Center, Yaling Yi, and John Englehardt for providing the ferret stool samples. We also thank A. Rueven, J. Bliska, N. Kordana, and K. Barrack for assistance with sample collection and processing, and L. Taub, C. Kestheley and Henry Dione with assisting the analysis. We also thank M. Martinez in the GDSC for preprocessing and linear modeling of the metagenomics data. This work was supported by the National Institutes of Health (NIH, NIH ES 033988-01A1) and Cystic Fibrosis Foundation (005912G223) to G.A.O., NIH T32HL134598 to M.M.C, R01AI164772, U01AI163063 and a subaward from the Dartmouth Cystic Fibrosis Research Center (P30DK117469; to M.S.S.), and NHLBI Federal contract #75N92024C00008 and Cystic Fibrosis Foundation #ENGELH21XX0 JFE. This work was also supported by NIH grant P20-GM130454.

