## Supplemental Tables 1-9 for "Profiling Bile Acids in the Stools of Humans and Animal Models of Cystic Fibrosis"

**Table S1. Statistical analysis for data shown in Figure 1**. To assess statistical significance, a mixed-effects linear model was applied to the log2 concentrations of bile acids for each functional Bile Acid Type, setting Genotype (NonCF and CF) as a fixed effect. The model accounted for repeated measures by treating Sample (or participant) as a random effect. Additionally, Batch was included as a fixed effect to control for variability resulting from multiple sample submissions. P-values were adjusted for multiple comparisons using FDR method. Red, bold text indicates $P < 0.05$.

| Bile Acid Type | Genotype p-value | Genotype p-value Adjusted | Batch p-value | Batch p-value Adjusted |
|---|---|---|---|---|
| Primary uBA | 0.1020 | 0.4331 | 0.3698 | 0.5177 |
| Secondary uBA | 0.1237 | 0.4331 | 0.9297 | 0.9297 |
| Primary cBA | 0.9685 | 0.9910 | 0.0001 | 0.0008 |
| Secondary cBA | 0.7758 | 0.9910 | 0.0153 | 0.0268 |
| Synthetic Intermediates & Atypical BAs | 0.9910 | 0.9910 | 0.0007 | 0.0025 |
| Secondary Metabolite | 0.3986 | 0.9300 | 0.4461 | 0.5205 |
| Hepatic Detox Products | 0.9015 | 0.9910 | 0.0146 | 0.0268 |

**Table S2. Statistical analysis for data shown in Figure S3-8**. Top 20 bile acids shown. To assess statistical significance, a linear model was applied to the log2 concentrations of each bile acid, setting Genotype (NonCF and CF) as a fixed effect. Batch effects were included as a fixed effect to control for variability arising from multiple sample submissions. P-values were adjusted for multiple comparisons using FDR method. Red, bold text indicates P < 0.05.

| Bile Acids | Genotype p-value | Genotype p-value Adjusted | Batch p-value | Batch p-value Adjusted |
|---|---|---|---|---|
| 7alfa OH 3 oxo 4 cholestenoic acid | 0.0006 | 0.0484 | 0.0249 | 0.0614 |
| Alloisolithocholic acid | 0.0048 | 0.2023 | 0.0027 | 0.0091 |
| Glycodeoxycholic acid 3 S | 0.0097 | 0.2284 | 0.4508 | 0.4918 |
| Taurolithocholic acid 3 S | 0.0109 | 0.2284 | 0.0000 | 0.0000 |
| Dehydrolithocholic acid | 0.0163 | 0.2731 | 0.3711 | 0.4270 |
| Glycochenodeoxycholic acid | 0.0206 | 0.2879 | 0.0726 | 0.1325 |
| alfa Muricholic acid | 0.0252 | 0.2890 | 0.1488 | 0.2192 |
| Isolithocholic acid | 0.0326 | 0.2890 | 0.3709 | 0.4270 |
| Glycodeoxycholic acid | 0.0329 | 0.2890 | 0.4765 | 0.5131 |
| Dioxolithocholic acid | 0.0386 | 0.2890 | 0.1149 | 0.1755 |
| Deoxycholic acid 3 G | 0.0404 | 0.2890 | 0.0000 | 0.0000 |
| Taurodehydrocholic acid | 0.0425 | 0.2890 | 0.0949 | 0.1505 |
| Lithocholic acid | 0.0447 | 0.2890 | 0.2992 | 0.3927 |
| Taurodeoxycholic acid | 0.0490 | 0.2943 | 0.3819 | 0.4335 |
| Deoxycholic acid 24 G | 0.0553 | 0.3099 | 0.0449 | 0.0972 |
| Glycoursodeoxycholic acid | 0.0629 | 0.3302 | 0.0878 | 0.1447 |
| Lithocholic acid 3 G | 0.0721 | 0.3561 | 0.0000 | 0.0001 |
| Lithocholic acid 3 S | 0.0802 | 0.3744 | 0.2796 | 0.3850 |
| Deoxycholic acid | 0.1002 | 0.4353 | 0.5594 | 0.5801 |
| Glycolithocholic acid | 0.1037 | 0.4353 | 0.0263 | 0.0631 |

**Table S3. Bile acid panel for targeted analysis of bile acids.**

| Targeted Panel for Continued Analysis | | | | |
|---|---|---|---|---|
| Bile Acid Type | Bile Acids | | | |
| Secondary Metabolite | Dehydro-LCA | 7-Keto-LCA | Allo-Iso-LCA | Apo-CA |
| | Hyo-DCA | UDCA | Iso-LCA | 3-Oxo-CA |
| | 7-Keto-DCA | 7,12-Di-Oxo-LCA | | |
| Primary uBA | CA | CDCA | Alpha-MCA | Beta-MCA |
| Secondary uBA | DCA | LCA | Omega-MCA | |
| Primary cBA | (G)-CDCA[-m] | | | |
| Hepatic Detox Product | LCA-3S[-f] | CA-3S[-h, -f] | | |
| Synthetic Intermediates & Atypical BAs | Allo-CA | | | |

[-h] BA is not detected in human samples
[-m] BA is not detected in mouse samples
[-f] BA is not detected in ferret samples

**Table S4. Targeted analysis for functional groups of bile acids from children with CF in Figure 2A**. To assess statistical significance, a mixed-effects linear model was applied to the log2 concentrations of bile acids for each functional Bile Acid Type, setting Genotype (NonCF and CF) as a fixed effect. The model accounted for repeated measures by treating Sample (or participant) as a random effect. P-values were adjusted for multiple comparisons using FDR method. Red, bold text indicates $P < 0.05$.

| Bile Acid Type | Model Type | Genotype p-value | Genotype p-value Adjusted |
|---|---|---|---|
| Primary uBA | Mixed | 0.0010 | 0.0061 |
| Secondary uBA | Mixed | 0.6869 | 0.8243 |
| Secondary Metabolites | Mixed | 0.5904 | 0.8243 |
| Synthetic Intermediates & Atypical BAs | Simple | 0.0025 | 0.0076 |
| Primary cBA | Simple | 0.8255 | 0.8255 |
| Hepatic Detox Product | Simple | 0.4304 | 0.8243 |

**Table S5. Targeted analysis of individual bile acids from children with CF for Figures S9 and S10**. To assess statistical significance, a linear model was applied to the log2 concentrations of each bile acid in the targeted panel, setting Genotype (NonCF and CF) as a fixed effect. P-values were adjusted for multiple comparisons using FDR method. Red, bold text indicates P < 0.05.

| Bile Acid | Genotype p-value | Genotype p-value Adjusted |
|---|---|---|
| Beta Muricholic Acid | 0.0009 | 0.0183 |
| Allocholic Acid | 0.0025 | 0.0253 |
| Omega-Muricholic Acid | 0.0114 | 0.0692 |
| Chenodeoxycholic Acid | 0.0189 | 0.0692 |
| 3-Oxocholic Acid | 0.0189 | 0.0692 |
| Cholic Acid | 0.0208 | 0.0692 |
| Dehydrolithocholic Acid | 0.0512 | 0.1418 |
| Hyodeoxycholic Acid | 0.0567 | 0.1418 |
| Alfa Muricholic Acid | 0.1019 | 0.2264 |
| Ursodeoxycholic Acid | 0.1240 | 0.2480 |
| Alloisoithochilc Acid | 0.1393 | 0.2532 |
| Lithocholic Acid | 0.1720 | 0.2866 |
| Dioxolithocholic acid | 0.2770 | 0.4261 |
| Isolithocholic Acid | 0.3171 | 0.4530 |
| Lithocholic Acid-3S | 0.4304 | 0.5738 |
| Deoxycholic Acid | 0.5619 | 0.6253 |
| 7-Keto-Lithocolic Acid | 0.5628 | 0.6253 |
| 7-Keto Deoxycholic Acid | 0.5628 | 0.6253 |
| (G)-Chenodeoxycholic Acid | 0.8255 | 0.8689 |
| Apocholic Acid | 0.9433 | 0.9433 |

**Table S6. Targeted analysis for functional groups of bile acids from mice stool presented in Figure 4A**. To assess statistical significance, a mixed-effects linear model was applied to the log2 concentrations of bile acids for each functional Bile Acid Type, setting Genotype (NonCF and CF) as a fixed effect. The model accounted for repeated measures by treating Sample (or mouse) as a random effect. P-values were adjusted for multiple comparisons using FDR method.

| Bile Acid Type | Genotype p-value | Genotype p-value Adjusted |
|---|---|---|
| Primary uBA | 0.5109 | 0.7752 |
| Secondary uBA | 0.1203 | 0.5283 |
| Secondary Metabolite | 0.8440 | 0.8440 |
| Hepatic Detox Product | 0.6202 | 0.7752 |
| Synthetic Intermediates & Atypical BAs | 0.2113 | 0.5283 |

**Table S7. Targeted analysis of individual bile acids from mice stool, presented in Figures S15 and S16**. To assess statistical significance, a linear model was applied to the log2 concentrations of each bile acid in the targeted panel, setting Genotype (NonCF and CF) as a fixed effect. P-values were adjusted for multiple comparisons using the FDR method. Red, bold text indicates P < 0.05.

| Bile Acid | Genotype p-value | Genotype p-value Adjusted |
|---|---|---|
| Dehydrolithocholic Acid | 0.0008 | 0.0159 |
| Deoxycholic Acid | 0.0062 | 0.0616 |
| Hyodeoxycholic Acid | 0.0095 | 0.0633 |
| Dioxolithocholic acid | 0.0259 | 0.1297 |
| Lithocholic Acid | 0.0353 | 0.1413 |
| Isolithocholic Acid | 0.0458 | 0.1526 |
| Apocholic Acid | 0.1181 | 0.3374 |
| Chenodeoxycholic Acid | 0.1531 | 0.3828 |
| Allocholic Acid | 0.2113 | 0.4473 |
| Alfa Muricholic Acid | 0.2236 | 0.4473 |
| Cholic Acid 3S | 0.2637 | 0.4557 |
| Omega-Muricholic Acid | 0.2734 | 0.4557 |
| 3-Oxocholic Acid | 0.2981 | 0.4587 |
| 7-Keto Deoxycholic Acid | 0.3699 | 0.5284 |
| Cholic Acid | 0.4474 | 0.5885 |
| 7-Keto-Lithocolic Acid | 0.4708 | 0.5885 |
| Ursodeoxycholic Acid | 0.5186 | 0.5963 |
| Alloisoithochilc Acid | 0.5367 | 0.5963 |
| Beta Muricholic Acid | 0.8108 | 0.8188 |
| Lithocholic Acid-3S | 0.8188 | 0.8188 |

**Table S8. Linear models for functional groups of BA measured from ferret stool presented in Figure 5A**. To determine statistical significance, a linear mixed-effects model was applied to the log2 transformed concentrations of bile acids for each functional bile acid type, setting Genotype (NonCF and CF) as a fixed effect. When needed, sample (or ferret) was set as a random effect to account for repeated measures. P-values were adjusted for multiple comparisons using the FDR method. Red, bold text indicates $P < 0.05$.

| Bile Acid Type | Genotype p-value | Genotype p-value Adjusted |
|---|---|---|
| Primary uBA | 0.0874 | 0.0874 |
| Secondary uBA | 0.0342 | 0.0427 |
| Secondary Metabolite | 0.0156 | 0.0259 |
| Primary cBA | 0.0068 | 0.0170 |
| Synthetic Intermediates & Atypical BAs | 0.0002 | 0.0010 |

**Table S9. Targeted analysis of individual bile acids from ferret stool, presented in Figure S18-19**. To assess statistical significance, a linear model was applied to the log2 concentrations of each bile acid in the targeted panel, setting Genotype (NonCF and CF) as a fixed effect. P-values were adjusted for multiple comparisons using the FDR method. Red, bold text indicates $P < 0.05$.

| Bile Acid | Genotype p-value | Genotype p-value Adjusted |
|---|---|---|
| 7-Keto Deoxycholic Acid | 0.0000 | 0.0003 |
| 7-Keto-Lithocolic Acid | 0.0002 | 0.0013 |
| Allocholic Acid | 0.0002 | 0.0013 |
| Dioxolithocholic acid | 0.0011 | 0.0050 |
| Apocholic Acid | 0.0056 | 0.0212 |
| (G)-Chenodeoxycholic Acid | 0.0068 | 0.0215 |
| Alfa Muricholic Acid | 0.0093 | 0.0254 |
| Omega-Muricholic Acid | 0.0126 | 0.0299 |
| Cholic Acid | 0.0143 | 0.0302 |
| Chenodeoxycholic Acid | 0.0321 | 0.0609 |
| Deoxycholic Acid | 0.0460 | 0.0795 |
| 3-Oxocholic Acid | 0.0742 | 0.1175 |
| Lithocholic Acid | 0.2091 | 0.3056 |
| Isolithocholic Acid | 0.2592 | 0.3517 |
| Dehydrolithocholic Acid | 0.2840 | 0.3597 |
| Beta Muricholic Acid | 0.3259 | 0.3642 |
| Hyodeoxycholic Acid | 0.3259 | 0.3642 |
| Ursodeoxycholic Acid | 0.7435 | 0.7848 |
| Alloisoithochilc Acid | 0.9677 | 0.9677 |
