## Supplemental Figures S1-20 for "Profiling Bile Acids in the Stools of Humans and Animal Models of Cystic Fibrosis"

A

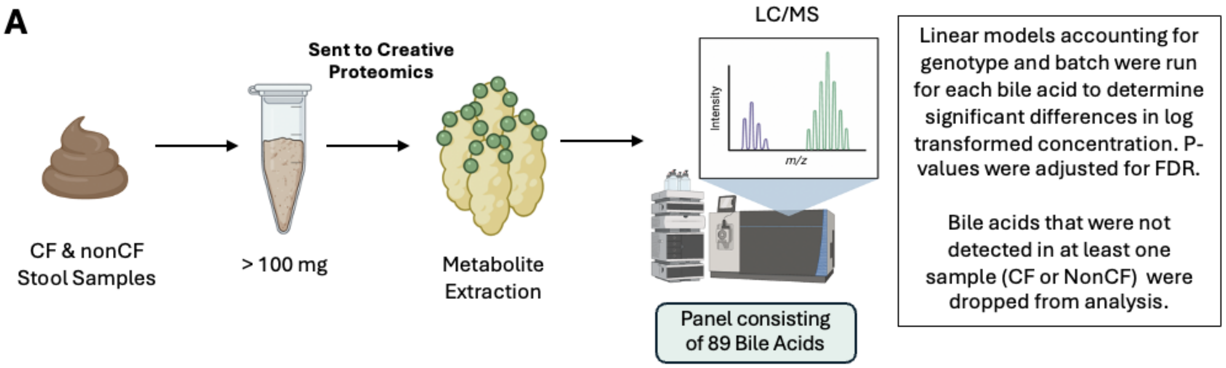


B

| | | | | | | |
|---|---|---|---|---|---|---|
| CA | LCA | 12-keto-CDCA | norUDCA | hyo-CA | (t)-DCA-3-SO4 | DCA-3-SO4 |
| CDCA | DCA | 12-keto-LCA | UCA | allo-CA | (t)-hyo-DCA-SO4 | hyo-DCA-24-gluc |
| MCA | omega-MCA | 3-oxo-CA | UDCA | (t)-hyo-CA | (t)-LCA-SO4 | hyo-DCA-3-gluc |
| Alpha-MCA | (g)-UDCA | 7-keto-DCA | (g)-hyo-DCA | hyo-DCA | (t)-UDCA-3-SO4 | hyo-DCA-3-SO4 |
| Beta-MCA | (g)-DCA | 7-keto-LCA | 3-beta-7-alpha-diOH-5-chol. | hyo-CA-3-SO4 | allo-CA-3-SO4 | iso-LCA-3-SO4 |
| (g)-CDCA | (g)-dehydro-CA | allo-iso-LCA | DHCA | (g)-allo-CA-3-SO4 | alpha-MCA-3-SO4 | LCA-24-gluc |
| (t)-CDCA | (g)-LCA | apo-CA | (g)-allo-CA | (g)-CA-3-SO4 | beta-MCA-3-SO4 | LCA-24-gluc |
| (t)-beta-MCA | (t)-DCA | dehydro-CA | 7-alpha-OH-3-oxo-4-chol. | (g)-CDCA-3-SO4 | CA-3-SO4 | LCA-3-SO4 |
| (g)-beta-MCA | (t)-dehydro-CA | dehydro-LCA | 6,7-di-keto-LCA | (g)-DCA-3-SO4 | CDCA-24-gluc | UDCA-24-gluc |
| (t)-alpha-MCA | (t)-LCA | di-oxo-LCA | 3-beta-OH-5-chol. | (g)-hyo-DCA-3-SO4 | CDCA-3-gluc | UDCA-3-gluc |
| (g)-alpha-MCA | (t)-omega-MCA | iso-DCA | (g)-hyo-CA | (g)-LCA-3-SO4 | CDCA-3-SO4 | UDCA-3-SO4 |
| (g)-CA | (t)-UCDA + (t)-hyo-DCA | norCA | (t)-allo-CA | (g)-UDCA-3-SO4 | DCA-24-gluc | |
| (t)-CA | iso-LCA | norDCA | THCA | (t)-CDCA-3-SO4 | DCA-3-gluc | |

**Legend**:

Primary uBA
Secondary Metabolites
Secondary uBA
Secondary cBA
Primary cBA
Hepatic Detox Products
Synthetic, Intermediates, and Atypical BAs

**Figure S1. Analysis using the 89 BA panel from Creative Proteomics. A.** Experimental workflow. **B.** BA panel from the untargeted Creative Proteomics colored by functional group. Legend of functional groups: Primary unconjugated BA, Primary conjugated BA, Secondary unconjugated BA, Secondary conjugated BA, Secondary metabolites, Hepatic detox products and Synthetic, Intermediates, and atypical BAs.

**A**

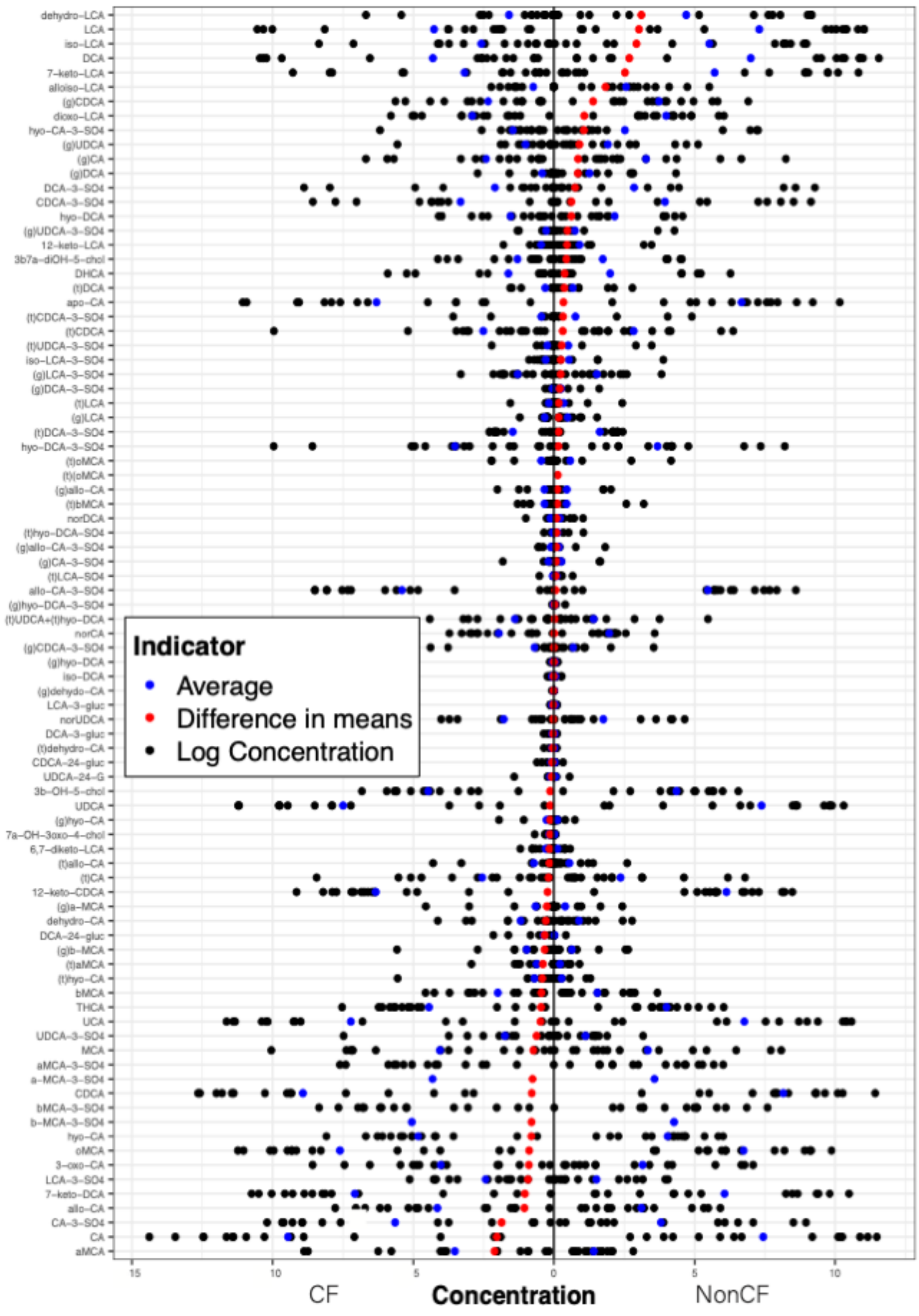

dehydro-LCA
LCA
iso-LCA
DCA
7-keto-LCA
alloiso-LCA
(g)CDCA
dioxo-LCA
hyo-CA-3-SO4
(g)UDCA
(g)CA
(g)DCA
DCA-3-SO4
CDCA-3-SO4
hyo-DCA
(g)UDCA-3-SO4
12-keto-LCA
3b7a-diOH-5-chol
DHCA
(t)DCA
apo-CA
(t)CDCA-3-SO4
(t)CDCA
(t)UDCA-3-SO4
iso-LCA-3-SO4
(g)LCA-3-SO4
(g)DCA-3-SO4
(t)LCA
(g)LCA
(t)DCA-3-SO4
hyo-DCA-3-SO4
(t)oMCA
(t)(oMCA
(g)allo-CA
(t)bMCA
norDCA
(t)hyo-DCA-SO4
(g)allo-CA-3-SO4
(g)CA-3-SO4
(t)LCA-SO4
allo-CA-3-SO4
(g)hyo-DCA-3-SO4
(t)UDCA+(t)hyo-DCA
norCA
(g)CDCA-3-SO4
(g)hyo-DCA
iso-DCA
(g)dehydo-CA
LCA-3-gluc
norUDCA
DCA-3-gluc
(t)dehydro-CA
CDCA-24-gluc
UDCA-24-G
3b-OH-5-chol
UDCA
(g)hyo-CA
7a-OH-3oxo-4-chol
6,7-diketo-LCA
(t)allo-CA
(t)CA
12-keto-CDCA
(g)a-MCA
dehydro-CA
DCA-24-gluc
(g)b-MCA
(t)aMCA
(t)hyo-CA
bMCA
THCA
UCA
UDCA-3-SO4
MCA
aMCA-3-SO4
a-MCA-3-SO4
CDCA
bMCA-3-SO4
b-MCA-3-SO4
hyo-CA
oMCA
3-oxo-CA
LCA-3-SO4
7-keto-DCA
allo-CA
CA-3-SO4
CA
aMCA
Indicator
Average
Difference in means
Log Concentration
15
10
5
0
5
10
CF
Concentration
NonCF

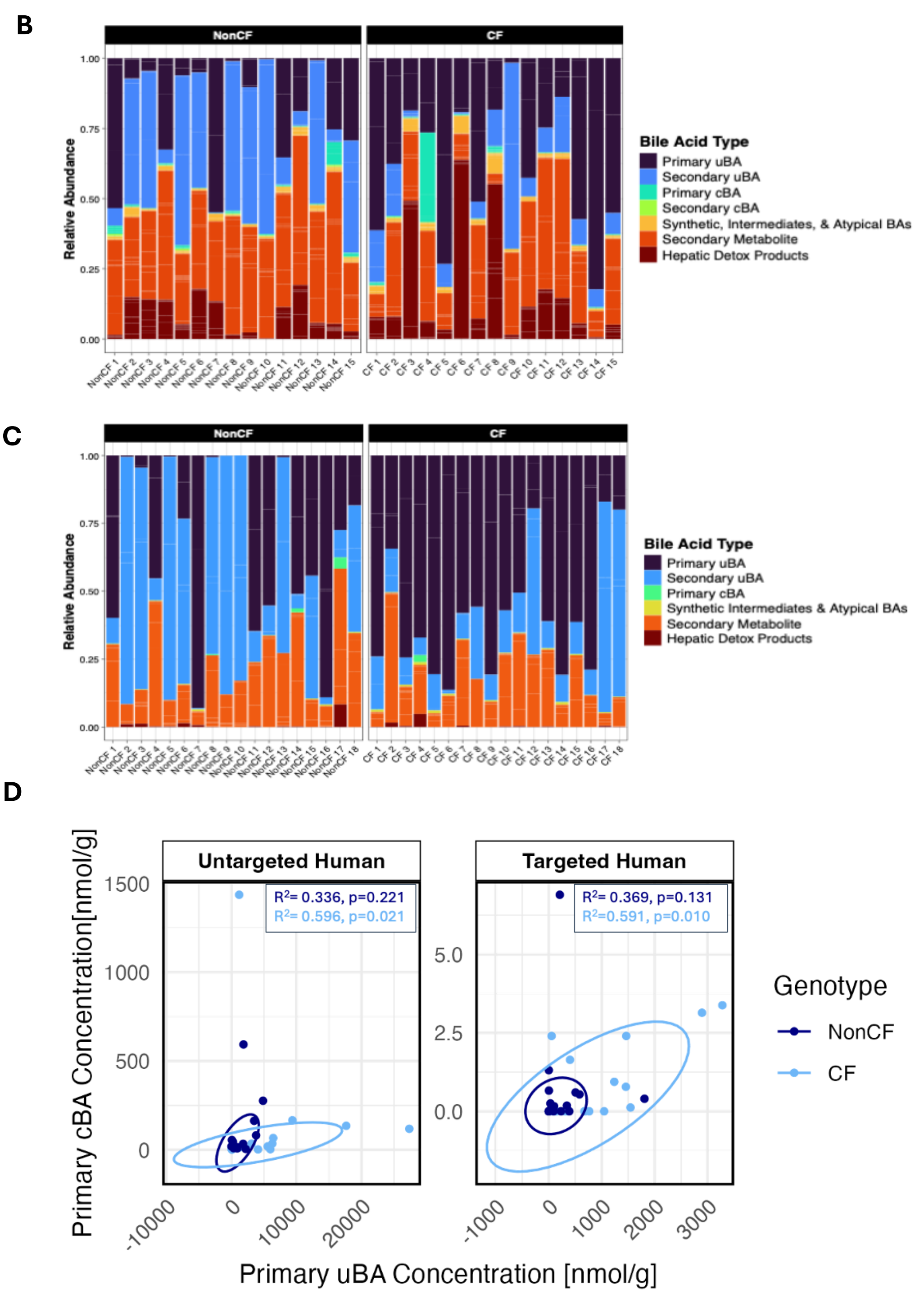


**Figure S2. BA analysis using the Creative Proteomics panel. A.** Difference in means of BA measured in the untargeted Creative Proteomics panel. The log concentration of BA in CF versus nonCF samples. The log2 transformed value for each sample is shown as a black dot, the average value for each cohort is shown as a blue dot and the difference in means is shown as a red dot for each BA. Legend is inset. **B**. Relative abundance of functional groups between genotypes (CF and nonCF) in individual human samples from the untargeted Creative Proteomics Panel. These are the data used to generate the graph in Figure 1C. **C**. Relative abundance of functional groups of the subset of BA analyzed (**Table S3**) between genotypes (CF and nonCF) in individual samples. These are the data used to generate the graph in Figure 2C. **D**. Relationship between primary cBA and primary uBA in humans. Scatter plot showing the relationship between primary unconjugated bile acid (Primary uBA) and primary conjugated bile acid (Primary cBA) concentrations (nmol/g) for each participant across the two metabolomics methods: A) Untargeted Human and B) Targeted Human. Data points are colored by genotype (NonCF in dark blue, CF in light blue). Confidence ellipses (95% confidence interval) illustrate the distribution and grouping of participants by genotype within each method. To determine the relationship between primary conjugated bile acids and primary unconjugated bile acids in this panel, bile acid concentrations were summed per sample by functional group (primary conjugated bile acids [cBA] and Primary unconjugated bile acids [uBA]) for each metabolomics method (Untargeted and Targeted) in human samples. To evaluate the significance and strength of the association between primary unconjugated and conjugated bile acids, Spearman correlation tests were performed for each metabolomics method and genotype group. The coefficient of determination ($R^2$) and corresponding p-values are reported using the same color scheme as the points (NonCF in dark blue, CF in light blue). Statistical analyses were performed in R.

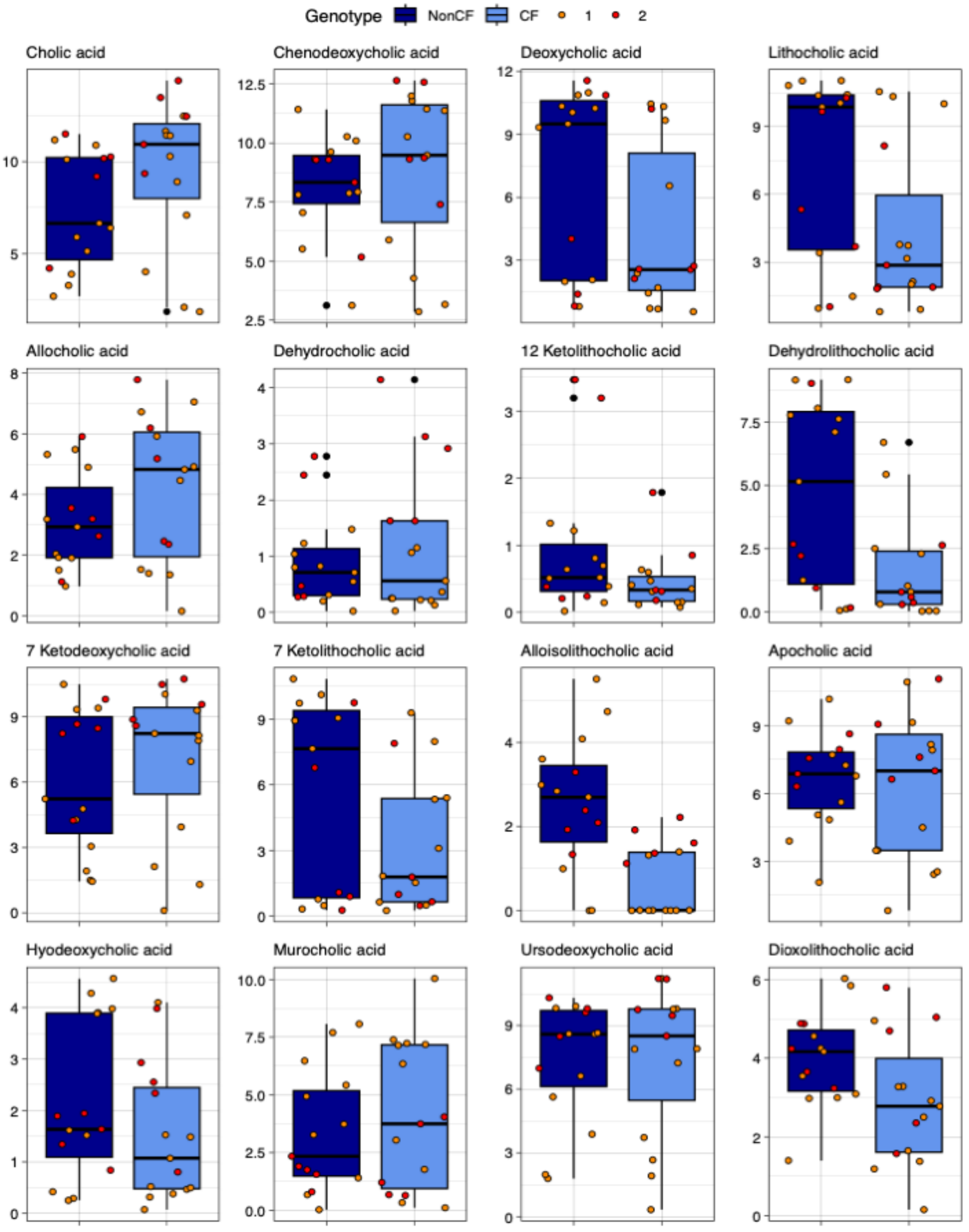


**Figure S3. Box plots comparing log2 transformed concentration for CF and nonCF human samples for BA in the Creative Proteomics Panel.** Colored by Genotype (CF and nonCF) and Batch (1,2). The corresponding statistical analyses are shown in **Table S2**.

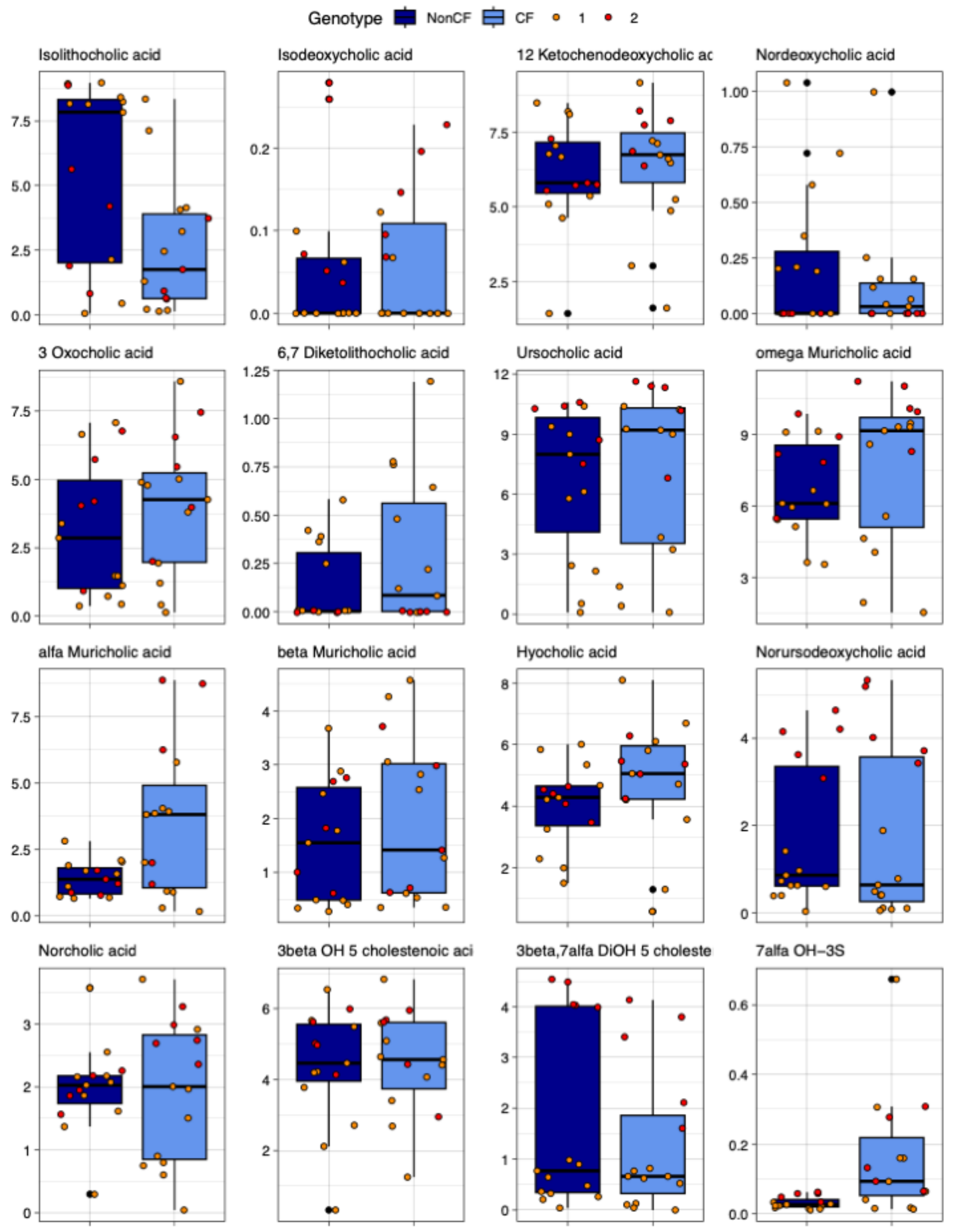


**Figure S4. Box plots comparing log2 transformed concentration for CF and nonCF human samples for BA in the Creative Proteomics Panel**. Colored by Genotype (CF and nonCF) and Batch (1,2). The corresponding statistical analyses are shown in **Table S2**.

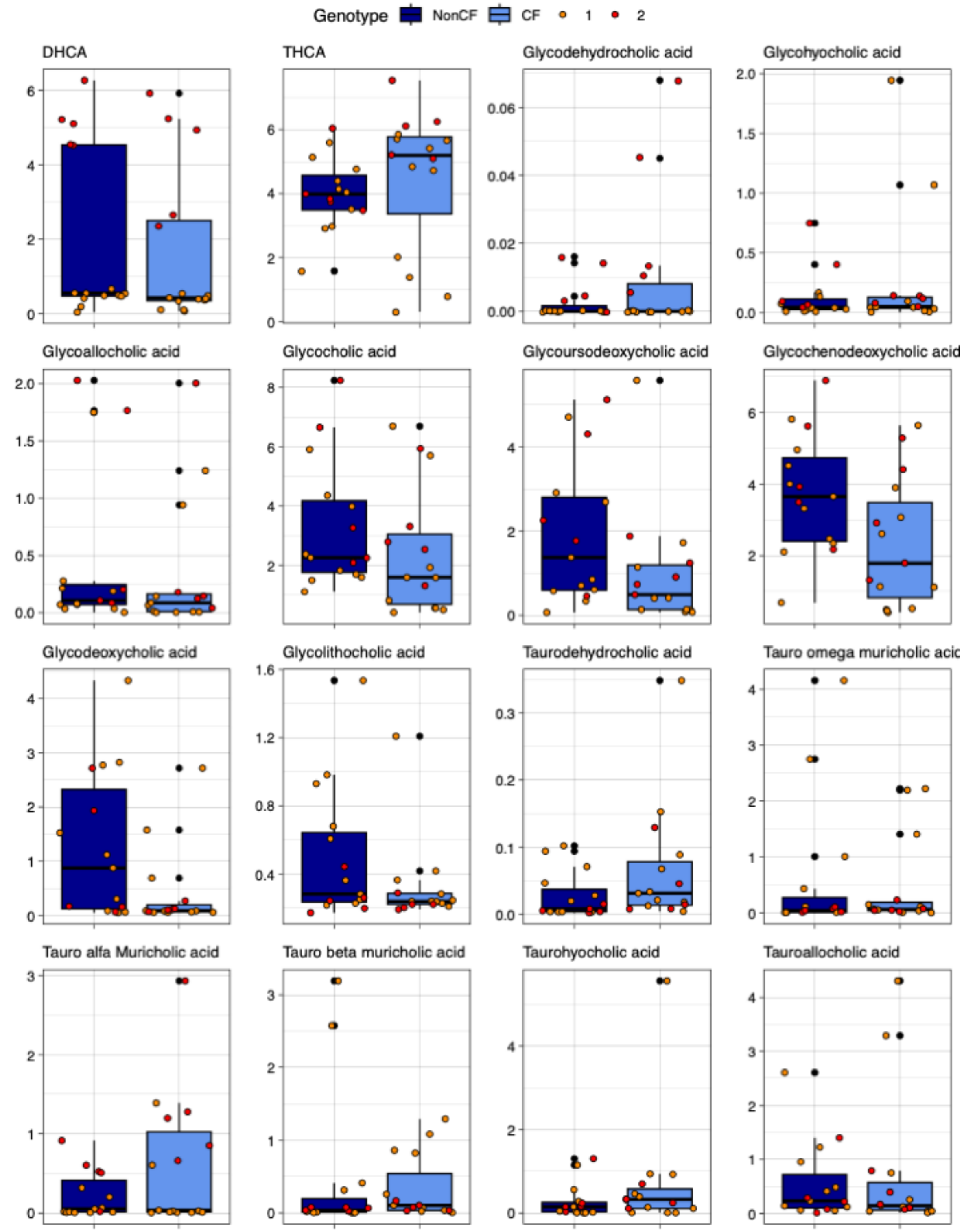


**Figure S5. Box plots comparing log2 transformed concentration for CF and nonCF human samples for BA in the Creative Proteomics Panel**. Colored by Genotype (CF and nonCF) and Batch (1,2). The corresponding statistical analyses are shown in **Table S2**.

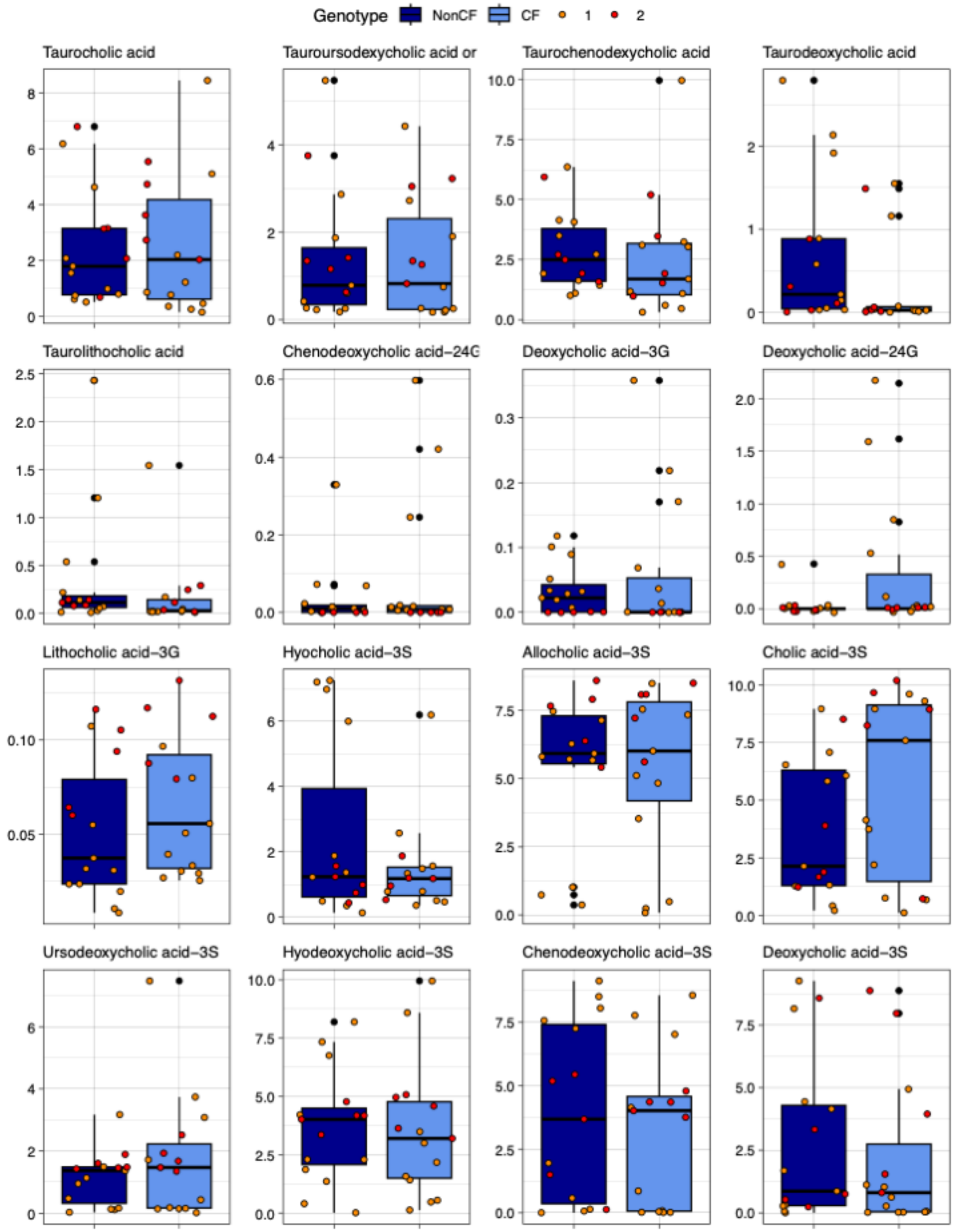


**Figure S6. Box plots comparing log2 transformed concentration for CF and nonCF human samples for BA in the Creative Proteomics Panel**. Colored by Genotype (CF and nonCF) and Batch (1,2). The corresponding statistical analyses are shown in **Table S2**.

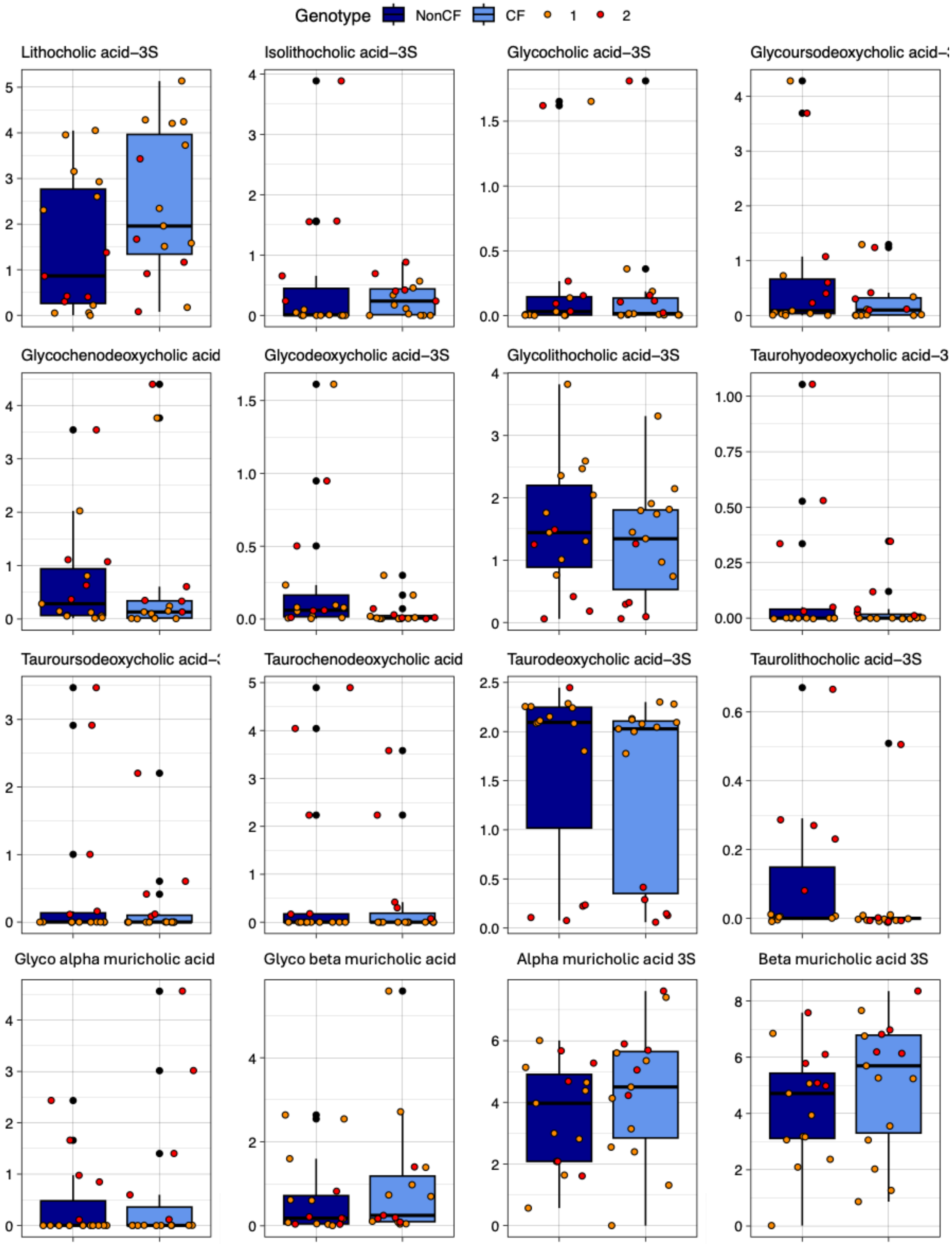


**Figure S7. Box plots comparing log2 transformed concentration for CF and nonCF human samples for BA in the Creative Proteomics Panel**. Colored by Genotype (CF and nonCF) and Batch (1,2). The corresponding statistical analyses are shown in **Table S2**.

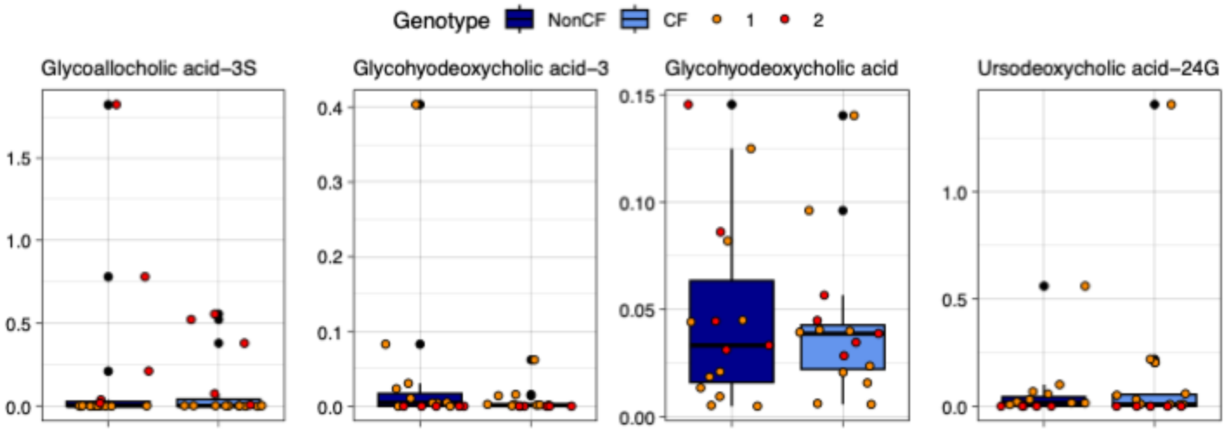


**Figure S8. Box plots comparing log2 transformed concentration for CF and nonCF human samples for BA in the Creative Proteomics Panel**. Colored by Genotype (CF and nonCF) and Batch (1,2). The corresponding statistical analyses are shown in **Table S2**.

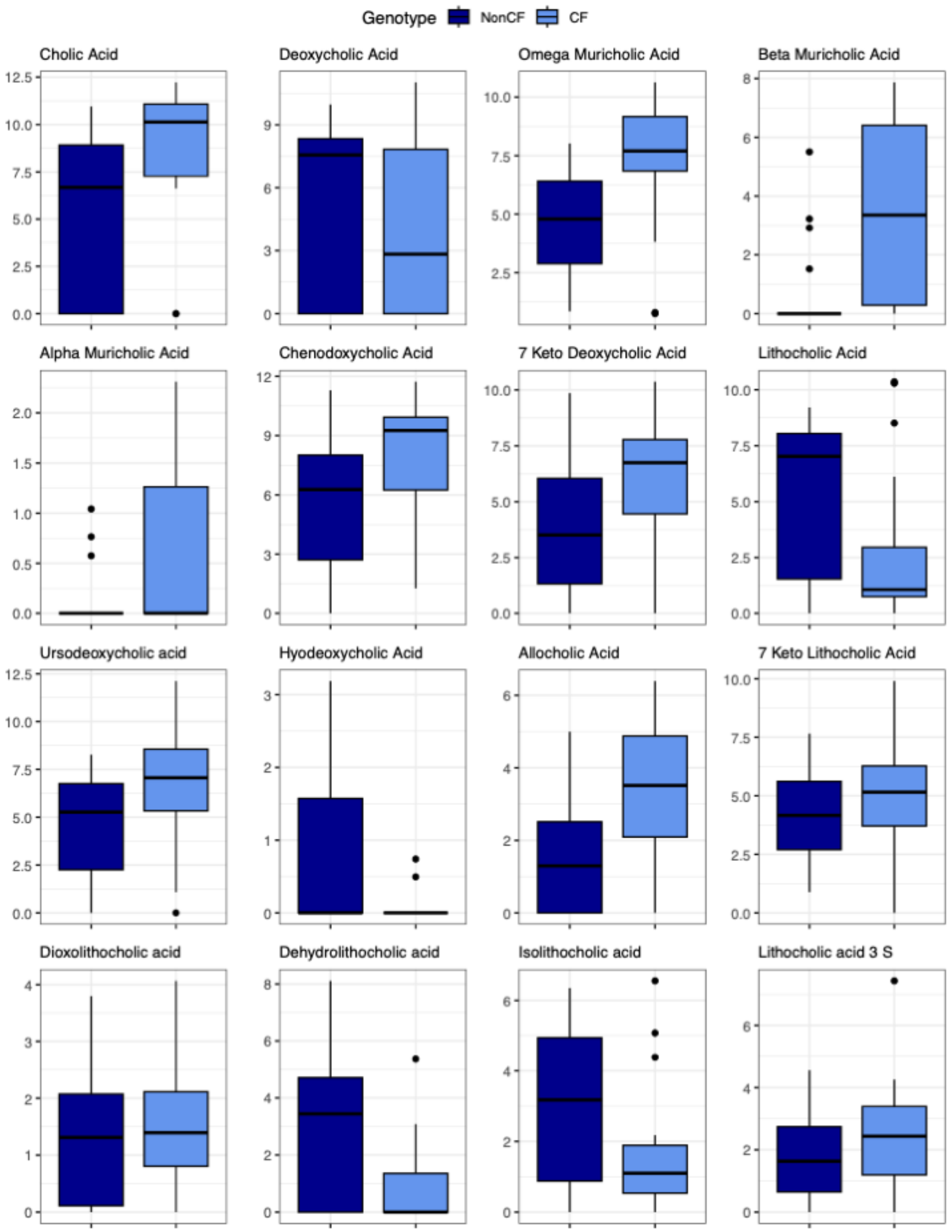


**Figure S9. Box plots comparing log2 transformed concentration of selected BA for CF and nonCF human samples**. Colored by Genotype (CF and nonCF). Samples were analyzed at the Mass Spectrometry and Metabolomics Core at Michigan State University. See the main text for details of the methods. The corresponding statistical analyses are presented in **Table S5**.

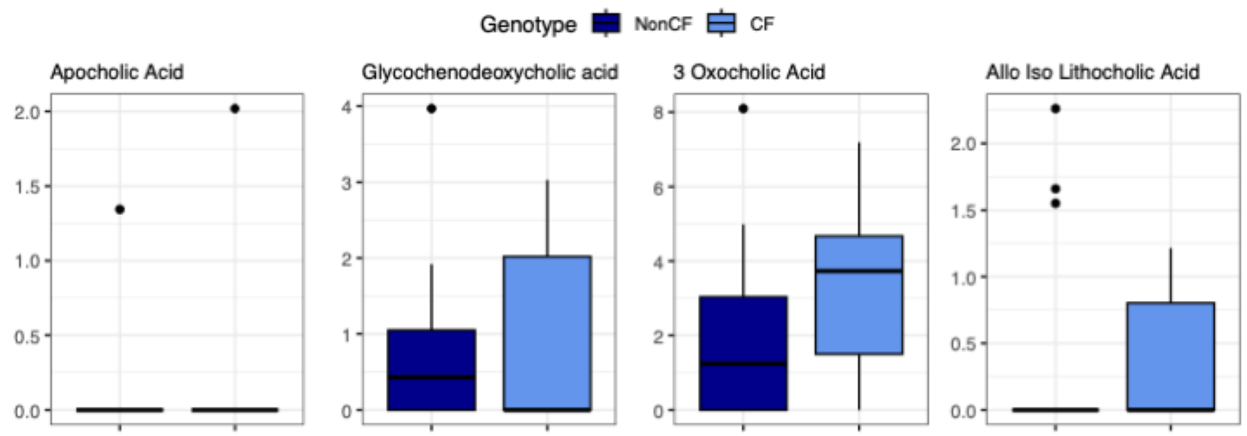


**Figure S10. Box plots comparing log2 transformed concentration of selected BA for CF and nonCF human samples**. Colored by Genotype (CF and nonCF). Samples were analyzed at the Mass Spectrometry and Metabolomics Core at Michigan State University. See the main text for details of the methods. The corresponding statistical analyses are presented in **Table S5**.

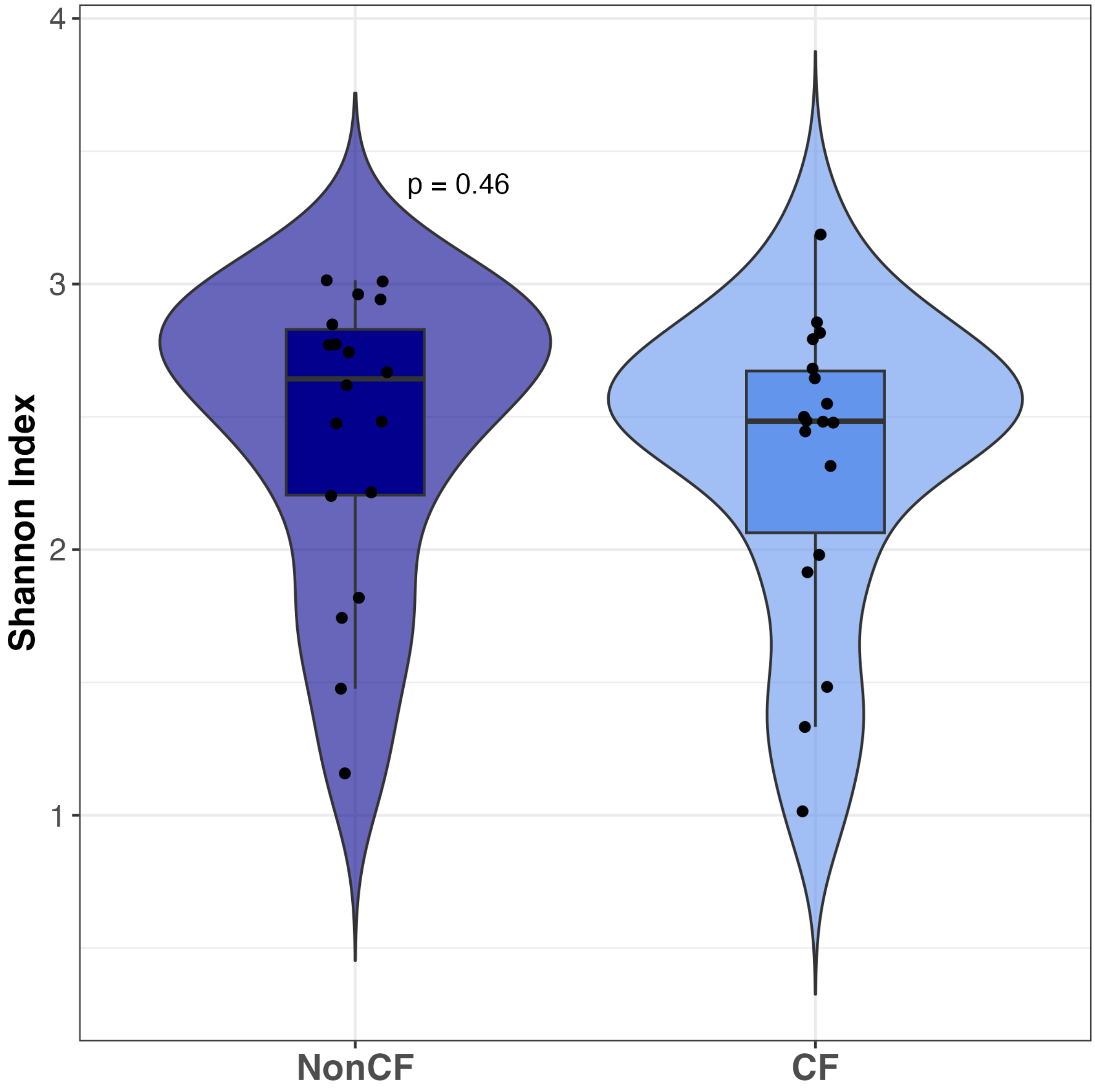


**Figure S11. Shannon Diversity.** Alpha diversity was calculated using the diversity function from vegan (v2.6.8). Relative abundances of taxa per sample (*$p_i$*) are used to calculate Shannon's H such that: $H = -\Sigma p_i * log(p_i)$. The Shannon index places an emphasis on species diversity (number of species) and evenness (how evenly species are distributed). Higher values of *H* indicate more diversity, reflecting either a higher number of taxa or a more even distribution of taxa within a sample. Data were analyzed using species abundance filtered for species with at least 10% prevalence across all samples. No significant difference between groups by Wilcox Test.

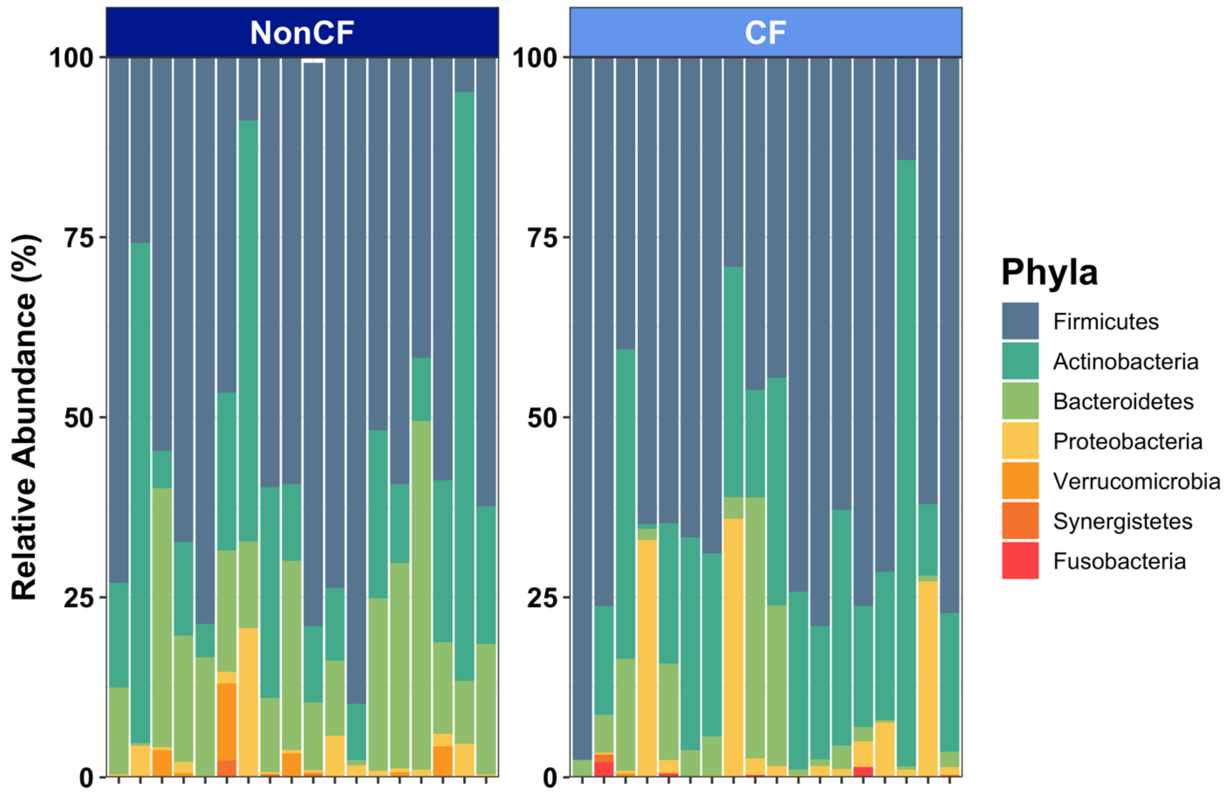


**Figure S12. Metaphlan taxonomic abundances between genotypes (CF and NonCF) in individual human samples from Creative Proteomics Panel.** Phylum level analysis using Metaphlan as described in the Materials and Methods. The legend indicates the phylum color.

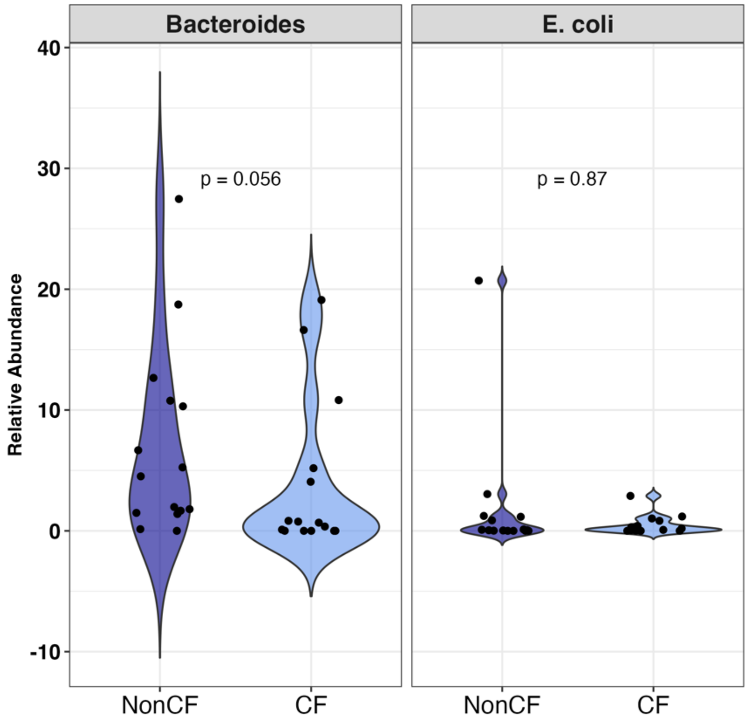


**Figure S13. Relative abundances of *Bacteroides* and *E. coli* between genotypes**. Violin plots illustrate the distribution of microbial relative abundances within each genotype group (CF vs NonCF) with individual data points overlaid. Statistical significance of differential abundance was assessed using the non-parametric Wilcox test. n = 15 per genotype.

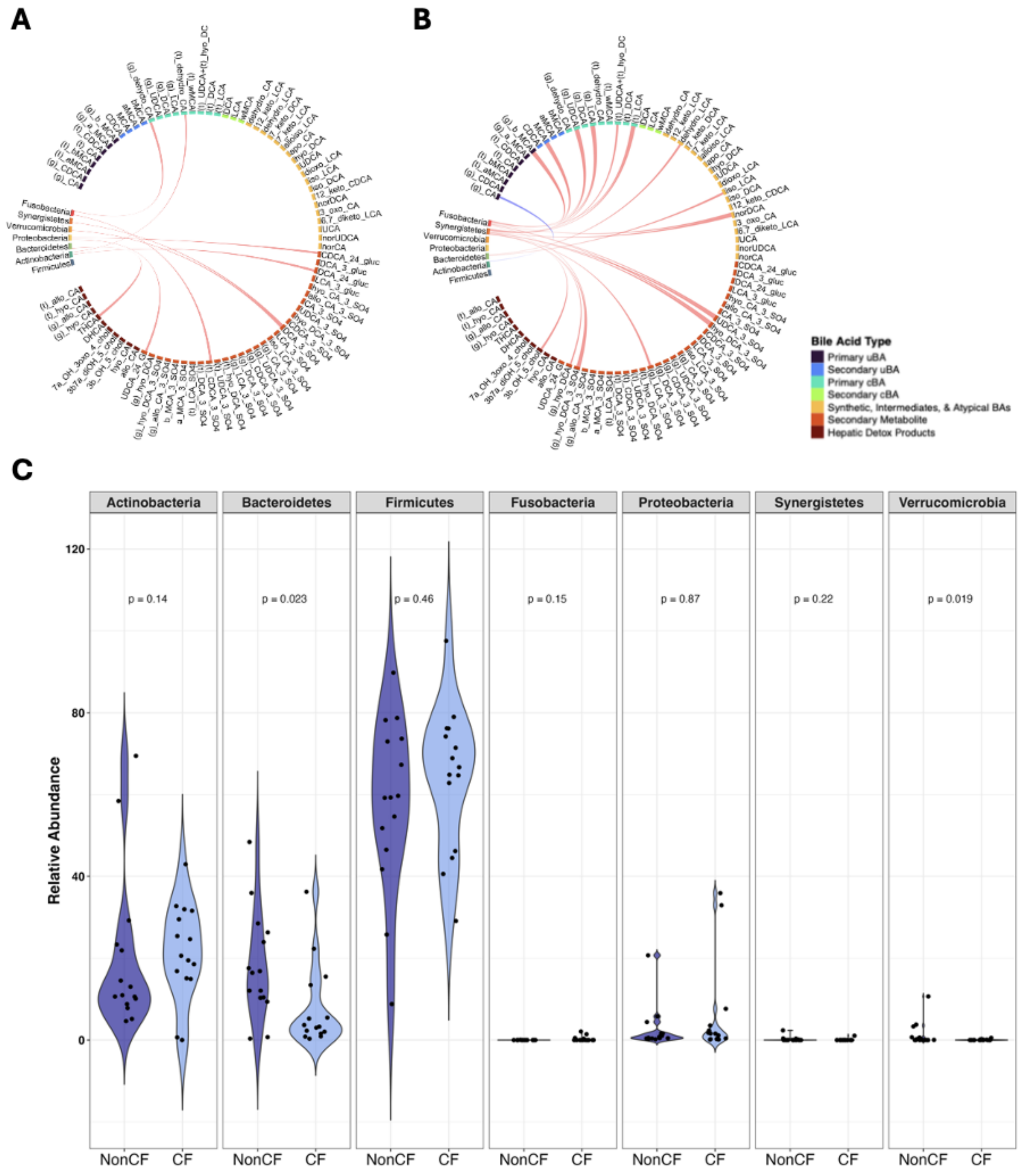


**Figure S14. Correlation plots of BA levels and microbes.** Circos plot assessing associations between BA and phyla for the (**A**) WT and (**B**) CF samples. Correlations are determined using a Pearson test, and a link is drawn if the correlation meets the threshold of 0.7 as follows: > 0.7 (red) or < -0.7 (blue). The class of BA is shown in the legend, and the color scheme of the phyla is consistent with **Figure S12**. **C.** Relative abundance of each phyla determined from the analysis of the metagenomes.

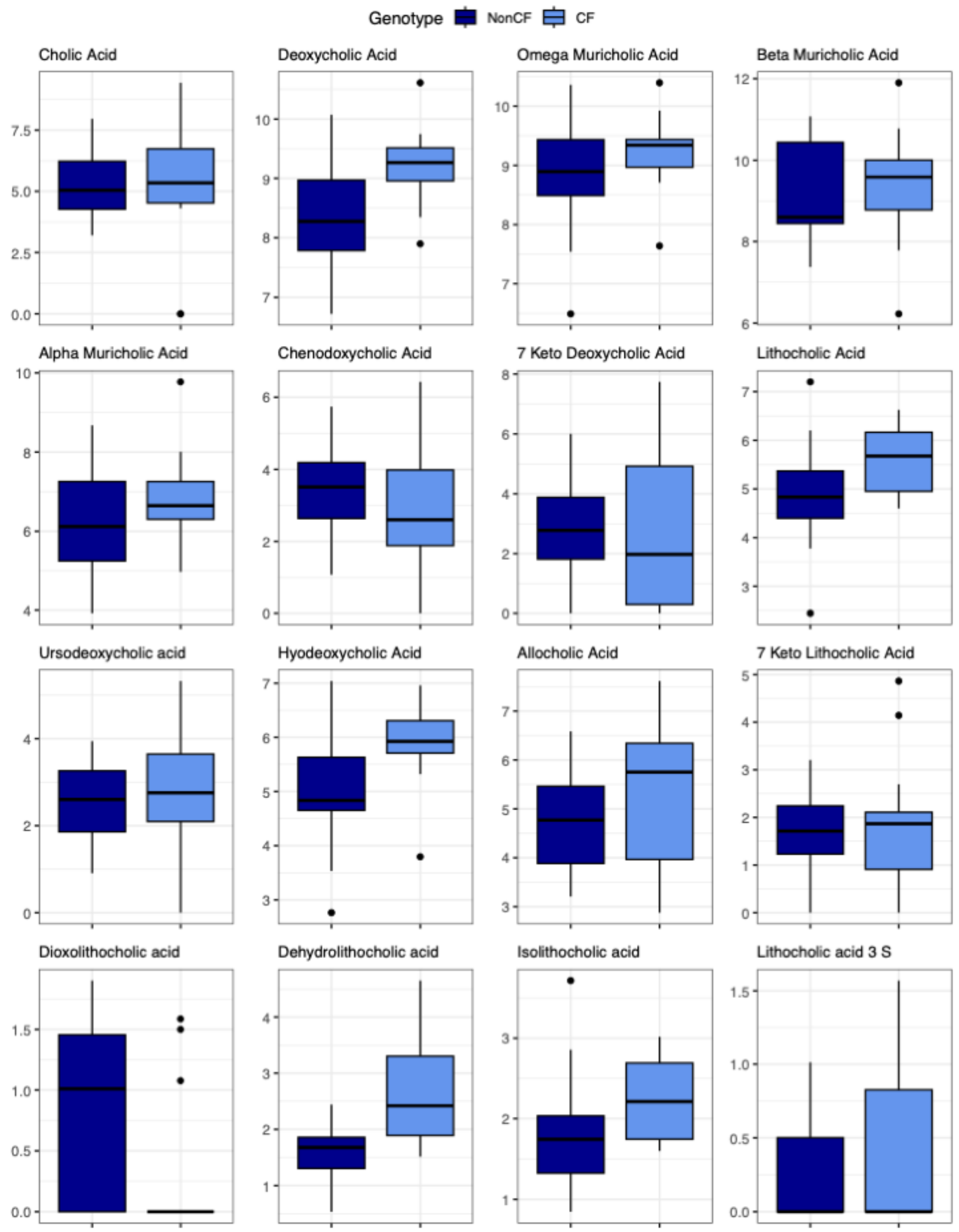


**Figure S15. Box plots comparing log2 transformed concentration of selected BA for CF and nonCF mouse samples**. Colored by Genotype (CF and nonCF). Samples were analyzed at the Mass Spectrometry and Metabolomics Core at Michigan State University. See the main text for details of the methods. The corresponding statistical analyses are presented in **Table S7**.

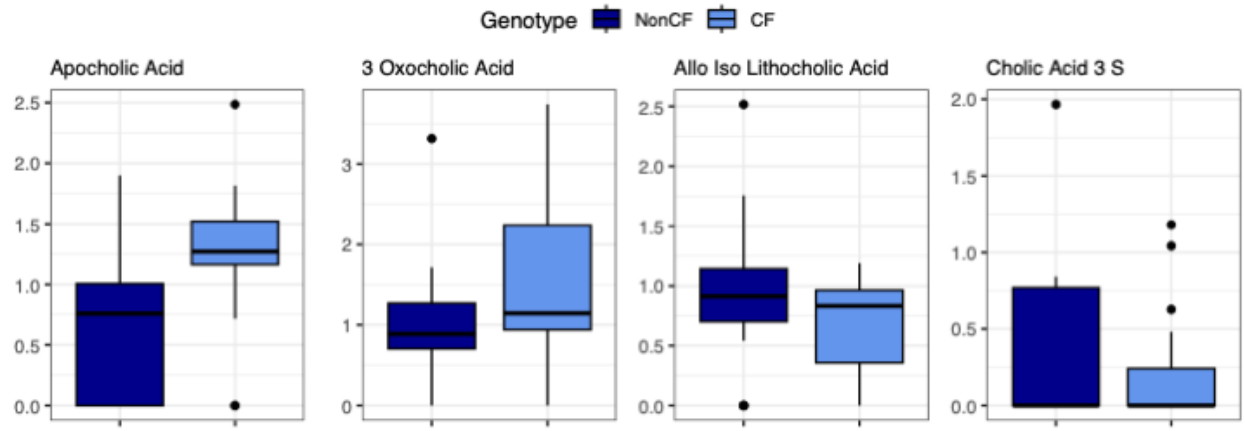


**Figure S16. Box plots comparing log2 transformed concentration of selected BA for CF and nonCF mouse samples**. Colored by Genotype (CF and nonCF). Samples were analyzed at the Mass Spectrometry and Metabolomics Core at Michigan State University. See the main text for details of the methods. The corresponding statistical analyses are presented in **Table S7**.

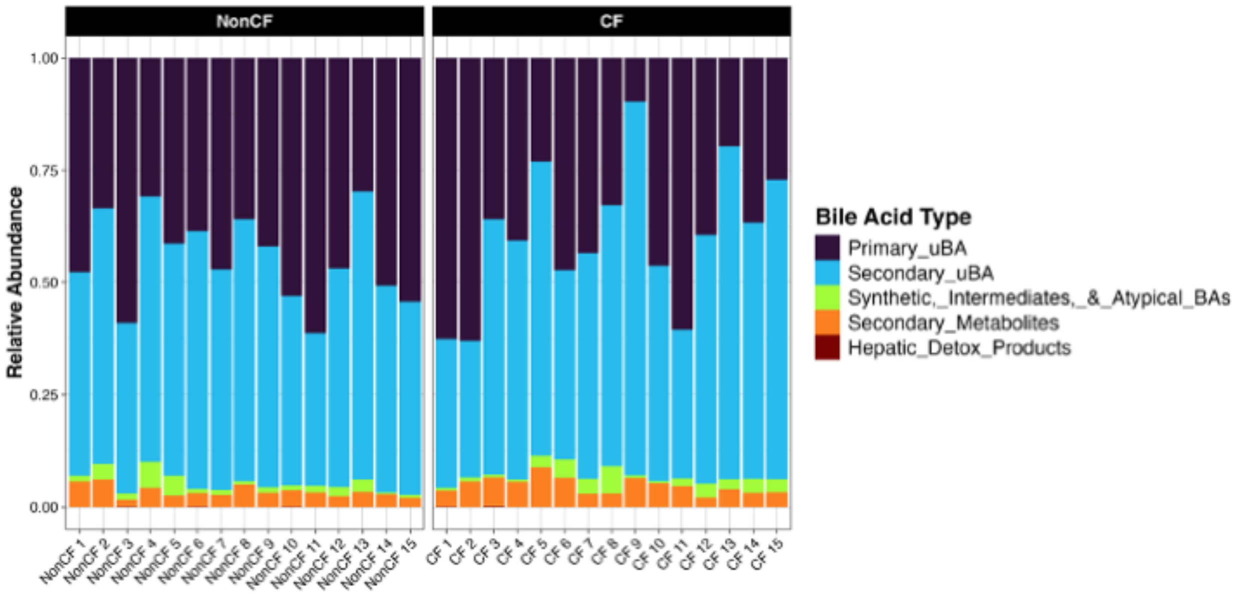


**Figure S17. Relative abundance of BA functional groups between genotypes (CF and nonCF) in individual mouse samples.** These are the data used to generate the graph in Figure 4C.

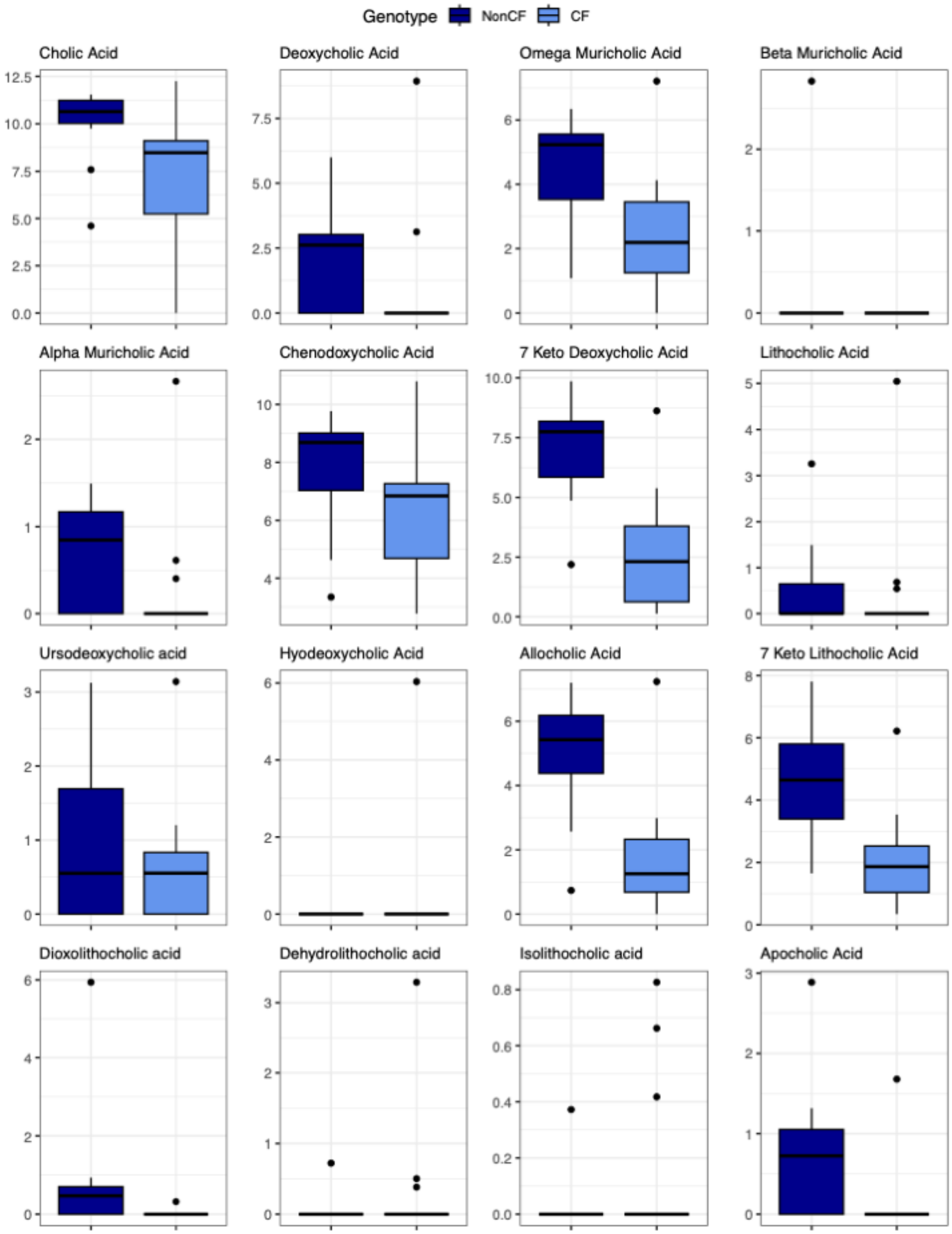


**Figure S18. Box plots comparing log2 transformed concentration of selected BA for CF and nonCF ferret samples**. Colored by Genotype (CF and NonCF). Samples were analyzed at the Mass Spectrometry and Metabolomics Core at Michigan State University. See the main text for details of the methods. The corresponding statistical analyses are presented in **Table S8**.

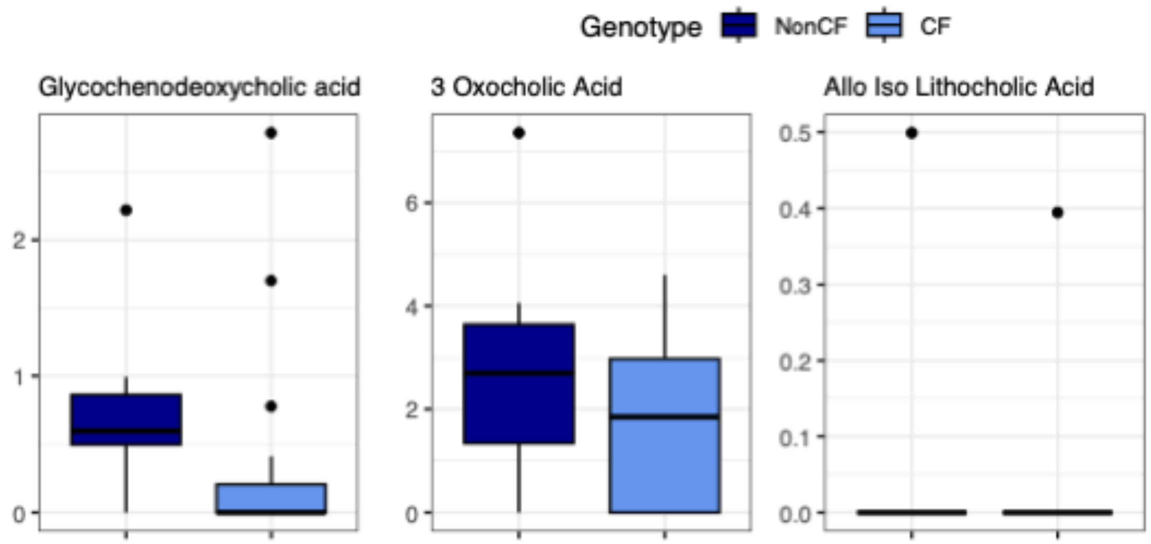


**Figure S19. Box plots comparing log2 transformed concentration of selected BA for CF and nonCF ferret samples**. Colored by Genotype (CF and NonCF). Samples were analyzed at the Mass Spectrometry and Metabolomics Core at Michigan State University. See the main text for details of the methods. The corresponding statistical analyses are presented in **Table S8**.

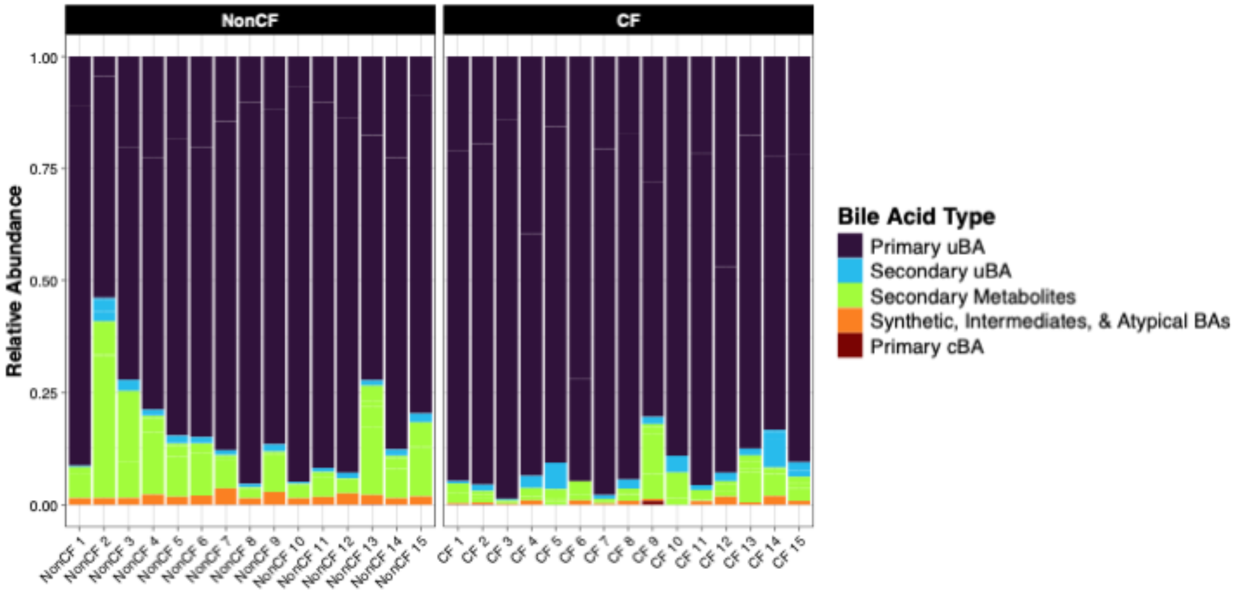


**Figure S20. Relative abundance of BA functional groups between genotypes (CF and nonCF) in individual ferret samples.** These are the data used to generate the graph in Figure 5C.
